# Ripened plant-based cheese analogs in Europe: nutritional and microbial profiles

**DOI:** 10.1101/2024.04.13.589336

**Authors:** Isabela Jaeger, Cecília R. Köhn, Joshua D. Evans, Jeverson Frazzon, Pierre Renault, Caroline Isabel Kothe

## Abstract

Plant-based cheese analogs have emerged as a novel global market trend driven by sustainability concerns for our planet. This study examines eleven soft ripened plant-based cheese analogs produced in Europe, primarily with bloomy rinds and cashew nuts as the main ingredient. First, we focused on exploring the macronutrients and salt content stated on the labels, as well a detailed fatty acid analysis of the samples. Compared to dairy cheeses, plant-based cheeses share similarities in lipid content, but their fatty acid profiles diverge significantly, with higher ratio of mono- and polyunsaturated fatty acids such as oleic and linoleic acids. We also investigated the microbiota of these analog products, employing a culture-dependent and -independent approaches. We identified a variety of microorganisms in the plant-based cheeses, with *Lactococcus lactis* and *Leuconostoc mesenteroides* being the dominant bacterial species, and *Geotrichum candidum* and *Penicillium camemberti* the dominant fungal species. Most of the species characterized are similar to those present in dairy cheeses, suggesting that they have been inoculated as culture starters to contribute to the sensorial acceptance of plant-based cheeses. However, we also identify several species that are possibly intrinsic to plant matrices or originate from the production environment, such as *Pediococcus pentosaceus* and *Enterococcus* spp. This coexistence of typical dairy-associated organisms with plant associated species highlights the potential microbial dynamics inherent in the production of plant-based cheese. These findings will contribute to a better understanding of plant-based cheese alternatives, enable the development of sustainable products, and pave the way for future research exploring the use of plant-based substrates in the production of cheese analogues.

## 1. Introduction

In recent years, plant-based diets have gained remarkable traction, driven by greater consumer awareness of nutritional, ethical, and cultural aspects (Pohjolainen et al., 2015). This shift is partly driven by concerns over the sustainability challenges posed by conventional animal-derived products and their limited capacity to meet the nutritional needs of a growing global population (Jeske et al., 2018; Karwacka et al., 2020).

Of particular interest, dairy products such as cheeses, yogurts, and butters hold an essential place in diets worldwide. However, growing evidence suggests that these products contribute substantially to environmental degradation, with dairy cheese emitting an average of 11 kg of CO_2_eq, compared to just 2 kg of CO_2_eq emitted by tofu, a plant-based cheese alternative (Poore & Nemecek, 2018). As we face the need to build sustainable food systems to accommodate the projected nine billion people by 2050 (FAO, 2018), exploring alternative production methods that incorporate plant-based ingredients is emerging as a promising avenue to align with evolving consumer preferences for animal-free alternatives.

The surge of veganism, coupled with the recognized health benefits associated with plant-based proteins, is a key driving factor behind the expansion of the analog cheese market. Nonetheless, there remains a need for the industry to enhance the functional and nutritional properties of these products and, at the same time, improve their stability (Kamath et al., 2022; McClements & Grossmann, 2022). One important aspect to consider for these novel alternatives is their fatty acid composition. As plant-based cheeses are often derived from nuts, they may exhibit richness in mono- and polyunsaturated fatty acids (Jardim et al., 2023), contrasting with dairy cheeses, which are predominantly composed of saturated fatty acids (Paszczyk et al., 2022). Understanding these disparities holds significance not only in nutritional and health contexts, where saturated fats have been linked to adverse cardiovascular effects (Zong et al., 2016), but also in optimizing techniques for producing palatable and stable cheese alternatives.

Additionally, the microbial composition of fermented food plays a pivotal role in shaping their sensory and nutritional attributes. Microorganisms involved in the fermentation process contribute to flavor development, texture enhancement, and nutritional enrichment of the final product (Siddiqui et al., 2023; Zhang et al., 2023). Moreover, the presence of beneficial strains in plant-based cheeses reinforces their potential health benefits, emphasizing the importance of characterizing the microbial dynamics inherent to these products. Despite the increasing availability of plant-based cheeses in the market, literature on this subject remains limited. Notably, studies on the microbiology of plant-based cheeses are still sparse, with some artisanal productions of fermented cashew nuts revealing predominant genera inherent to the matrix, including *Lactococcus, Leuconostoc, Pediococcus* and *Weissella* (Chen et al., 2020; Tabanelli et al., 2018). Additionally, one of these studies has highlighted the potential allergen mitigation benefits of fermenting cashew nut-based products (Chen et al., 2020), showing the health advantages associated with fermentation.

In this context, our study aims to fill this gap by characterizing the fatty acid profiles and microbial communities present in plant-based cheese analogs collected from the European market and comparing them with dairy cheeses. It is noteworthy that while the term “plant-based cheese” is used in this article, it is essential to acknowledge that the European Union has prohibited dairy-related terminologies for plant-based product names (Court of Justice of the European Union, 2017).

To the best of our knowledge, this study represents the first attempt to collect samples of ripened plant-based cheeses from the market for comprehensive analysis of their fatty acid profiles and microbiota. The data collected here is promising not only for advancing research into plant-based cheese analogues, but also for providing valuable information for the development of novel food products.

## 2. Materials and Methods

### 2.1 Sampling and nutritional label profiles

A comprehensive search was conducted to identify ripened plant-based cheese products available across Europe. This search encompassed various sources, including websites specializing in plant-based products, magazine articles, and YouTube channels dedicated to discussions about plant-based cheeses. The samples selected for this study were deliberately chosen to include ripened alternatives aged for several weeks, rather than fresh products.

Before conducting some nutritional measurements, we compiled the nutritional information presented on the labels of the collected plant-based cheeses. This included examining the content of lipids, saturated fatty acids, carbohydrates, proteins, and salt, which were compared with existing data in the literature for dairy cheeses.

### 2.2 Protein, ash and fatty acids analyses

Analysis of protein, ash and fatty acids was performed with a homogeneous mixture of core and rind samples, in duplicate. The protein content of the samples was estimated using the Kjeldahl method using a correction factor of 5.30 with a digestion apparatus (Novatecnica NT415, Piracicaba, São Paulo, Brazil). The ash content was determined in a muffle furnace (Elektro Therm Linn, 312.6 SO LM1729, Germany) set to 550 °C. The fatty acid methyl esters (FAMEs) were determined by gas chromatography (GC Model 2010; Shimadzu, Kyoto, Japan) using fused silica column (Rtx-Wax, Restek, 30 m × 0.25 mm × 0.25 μm) (Joseph & Ackman, 1992) and the operation conditions were described by David et al. (2005). The fatty-acids quantification was conducted in accordance with the American Oil Chemists’ Society method (AOCS, 2003).

### 2.3 Microbiology

Using a sterile knife, the samples were separated into rind (1 g) and core (5 g) and added to 9 mL and 45 mL of physiological saline solution (0.9% NaCl), respectively. Subsequently, they were homogenized using the Lab Blender Stomacher® equipment (model 400) for 60 seconds at maximum speed. The homogenized content was used for culture-dependent and -independent analyses.

#### 2.3.1 Culture-dependent analyses

The viable lactic acid bacteria counts in the collected samples (rind and core) were estimated using M17 and De Man, Rogosa, and Sharpe (MRS) media. Total mesophilic counts were estimated using Brain Heart Infusion (BHI) media, and fungal counts using Yeast Extract Peptone Dextrose (YEPD) media. In order to prevent fungal growth, Amphotericin B (Sigma-Aldrich, St. Louis, MO, USA) was added at a final concentration of 20 mg/mL (50 mg/mL stock solution of Alfa Aeser™ Amphotericin B from *Streptomyces nodosus* in DMSO) to the M17, MRS, and BHI culture media. In the YEPD medium, 100 µg of ampicillin and 40 µg of chloramphenicol were added per mL of medium to inhibit bacterial growth.

For fungal counting on rinds, the samples were diluted from 10^-3^ to 10^-6^, cultured in the YEPD medium, and incubated at 25 °C for 48-72 hours. For bacterial counting, the core and rind samples were cultured in BHI, MRS, and M17 media and incubated at 30 °C under anaerobic and aerobic conditions, respectively. To create an oxygen-free environment, an anaerobic jar was used with GENbox sachets (bioMérieux, France).

For the counting and characterization of isolates from each sample, we selected plates containing 20 to 200 clones. The selected clones were collected using a sterile loop and mixed in a tube containing 300 μL of milliQ water, 100 mg of 0.1 mm diameter zirconia beads, and 10 mg of 0.5 mm diameter beads (Sigma, St. Louis, MO, USA). The tube was vigorously shaken using the Bead Beater equipment (FastPrep-24, MP Biomedicals Europe, Illkirch, France) for 20 seconds at 6.5 m/s. The supernatant from this lysis step was used directly for DNA amplification.

Species assignment was performed based on the sequencing of the 16S rRNA and 26S rRNA genes. The 16S rRNA gene was amplified using the primers 27-F (5’-AGAGTTTGATCATGGCTCA-3’) and 1492-R (5’-TACGGTTACCTTGTTTACGACTT-3’) (Acinas et al., 2004). The D1/D2 region of the 26S rRNA gene was amplified and sequenced using the primers NL-1 (5’-GCATATCAATAAGCGGAAAAG-3’) and NL-4 (5’-GGGTCCGTGTTTCAAGACGG-3’) (Kurtzman & Robnett, 1997). The PCR conditions for 16S rRNA were 1 min at 94 °C, 30 cycles of denaturation (1 min at 94 °C), annealing (0.5 min at 56 °C), and extension (1.5 min at 72 °C), followed by a final elongation (5 min at 72 °C). For the amplification of D1/D2 region, we used slightly different conditions: 2.5 min at 94 °C, 35 cycles of denaturation (1 min at 94 °C), annealing (1.5 min at 52 °C), and extension (2 min at 68 °C), followed by a final elongation (5 min at 68 °C). The DNA amplifications were separated on a 0.8% agarose gel. The PCR products were purified using ExoSAP-IT (Thermo Fisher Scientific, Waltham, MA, USA) and sent for sequencing at Eurofins Genomics, Ebersberg, Germany. The sequences were analyzed using the NCBI BLAST tool (Camacho et al., 2009) to obtain a taxonomic classification for each isolate.

#### 2.3.2 Culture-independent analyses

For DNA extraction 4 mL of the samples from Stomacher were collected and centrifuged (Sigma 1-15K Centrifuge) at 13,000 rpm at 4 °C for 3 minutes. Subsequently, DNA from the pellets was extracted using the DNeasy PowerFood Microbial Kit (Qiagen), following the manufacturer’s protocol.

The amplification of the V4 region of the 16S rRNA gene was performed using the 515-F and 806-R primers (Caporaso et al., 2011). The amplification procedure was carried out in a 25 μL volume, comprising genomic DNA (12.5 ng), 1.0 mM MgCl_2_, 0.5 μM of each primer, 0.2 mM dNTPs, 1X PCR Buffer, and 2U Platinum Taq DNA polymerase (Life Technologies). The amplification conditions for the 16S rRNA gene comprised an initial denaturation at 94 °C for 2 min, followed by 30 cycles of denaturation at 94 °C for 45 s, annealing at 55 °C for 45 s, and extension at 72 °C for 5 min. Similarly, the amplification of the Internal Transcribed Sequence (ITS) region of fungal rRNA was performed using the ITS1 and ITS2 primers (White et al., 1990). This amplification, conducted in a 25 μL volume, involved the inclusion of genomic DNA (12.5 ng), 2.5 mM MgCl_2_, 0.16 μM of each primer, 0.2 mM dNTPs, 1X PCR Buffer, and 1U Platinum Taq DNA polymerase (Life Technologies). The amplification conditions for ITS comprised an initial denaturation at 95 °C for 5 min, followed by 30 cycles of denaturation at 95 °C for 45 s, annealing at 56 °C for 45 s, and extension at 72 °C for 10 min. Subsequently, amplicons resulting from the PCR reactions were purified using Agencourt AMPure XP beads, following the guidelines of the manufacturer. DNA library indexing was performed following the instructions provided by Illumina Inc. (San Diego, California, USA). The sequencing process was executed utilizing an Illumina MiSeq System equipped with a v2 500-cycles kit.

The quality of the raw data was evaluated with FastQC (Wingett & Andrews, 2018). After, the sequences were imported into the FROGS pipeline (Escudié et al., 2018) to obtain the Operational Taxonomic Units (OTUs) and were filtered by length (150–500 bp). Subsequently, they were pooled into OTUs with SWARM (Mahé et al., 2014) with the aggregation distance cluster of 1. Chimeras were removed with VSEARCH (Rognes et al., 2016) and OTUs with abundances of less than 0.005% of sequences were filtered out, as recommended by Bokulich et al. (2013). The OTUs were affiliated with SILVA 132 SSU databases (Quast et al., 2013) for bacteria and UNITE 8.2 for fungi (https://unite.ut.ee/). Taxonomic composition and beta-diversity analyses were performed in R Studio v.3.6.1 using the phyloseq and ggplot2 packages v.1.30.0 (McMurdie & Holmes, 2013; Poirier et al., 2018).

Raw sequences of amplification of 16S rRNA gene and ITS are deposited on the European Nucleotide Archive (ENA) under the BioProject ID PRJNA1020213.

## 3. Results & Discussion

### 3.1 Sample macroscopic observations

Our research revealed that most plant-based ripened cheese analogs available on the market are crafted primarily from cashew nuts, and usually as camembert-style cheeses. In this study, we collected 11 plant-based soft cheese analogs, mainly with bloomy rinds, produced with cashew nuts, from France and the Netherlands (**Fig. 1A**). To diversify our sample set, we also included two cheese analogs made with almond milk as the primary ingredient (C10 and C11), as well as variations such as a sample using blue cheese technology (C8), and samples with natural and washed rinds (C2, C7 and C8). These samples represent the majority of ripened plant-based cheeses currently produced in the European market. Information about the starter cultures employed in production is usually absent on the labels; except for samples 5 and 6, which specify the use of *P. camemberti* (**Table S1**), the white mold typically used in dairy bloomy rind cheeses.

**Fig. 1.**
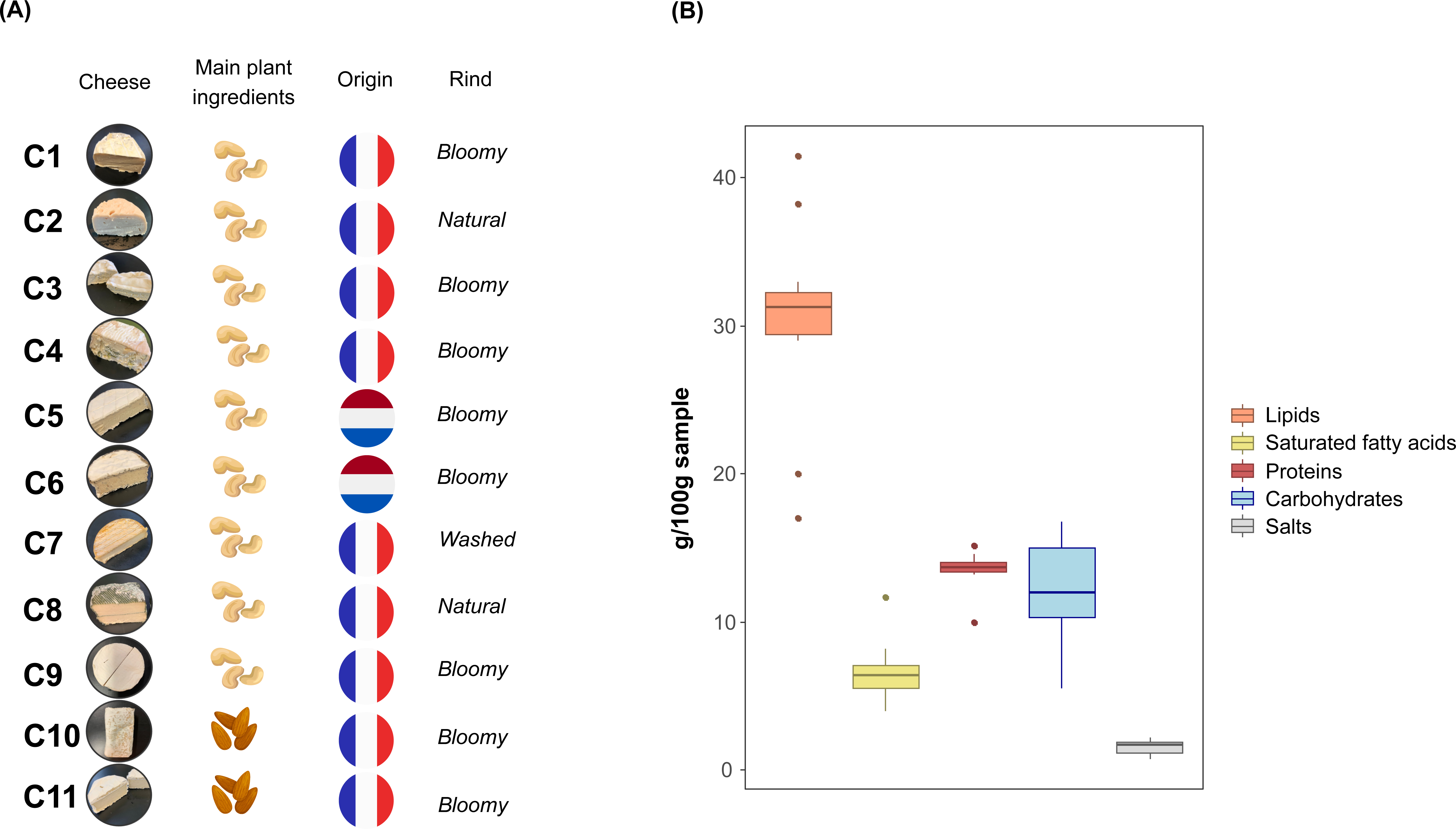
Ripened plant-based cheese analogs collected in this study with respective metadata (A) and nutritional composition based on label analysis (B).

Based on the labeling data, we analyzed the nutrient facts of our sample set. First, a significant amount of lipids was observed in the collected plant-based cheeses (30.18±6.95%, **Fig. 1B**). These fat values are similar to those found in Camembert and Brie dairy cheeses (Adamska et al., 2017; Bae et al., 2020). However, an important difference exists in the composition of the fatty acids. While saturated fatty acids represent 22% of total lipids in plant-based cheese, dairy cheeses have values three times higher. Saturated fatty acids, which are generally solid at room temperature, play a role in binding proteins and retaining moisture within the cheese paste, contributing to its cohesive structure, firmness and stability (Bugaud et al., 2001; Fox et al., 2017).

The protein content indicated on the labels averaged about 13.27±1.70%, which is statistically similar to our analytical measurements (11.94±1.58%, p > 0.05, **Table S2**). The protein content of the plant-based samples resembled that of Chinese *sufu*, a cheese-like product made from soy (Li & Wang, 2012). However, dairy cheeses typically exhibit higher protein content of about 15-20% (Batty et al., 2019; Li & Wang, 2012). Dairy milk contains casein and whey as proteins, while cashew nuts contain globulins, albumins, and glutelin (Liu et al., 2018). These distinctions in protein structures influence the processing techniques employed in the production of plant-based products. In traditional dairy cheese production, caseins from dairy milk are normally coagulated using rennet, a complex of enzymes. The absence of caseins in plant-based matrices requires alternative coagulation agents, such as calcium salts (Chen et al., 2021). Alternatively, another approach involves soaking the plant substrate in water, followed by blending the mixture to form a paste, which is subsequently fermented (Chen et al., 2020).

Carbohydrate content is on average 12.01±3.64% (**Fig. 1B**) in the plant-based cheese analogs from this study. This macronutrient can often be present at high levels in plant matrices, which represents a major challenge in creating plant cheeses. For example, cashew nuts, the main ingredient in the plant-based cheeses analyzed in this study, contain relatively higher levels of carbohydrates (>10%), predominantly composed of starch, which serve as an energy reserve polysaccharide in plants (Brufau et al., 2006; Chen et al., 2022). In contrast, dairy cheese contains only about 2% of lactose as primary carbohydrate (Cebeci et al., 2020), which is converted ultimately into lactic acid by lactic acid bacteria during the cheese-making process.

Salt plays an important role in preserving and enhancing flavor profiles in fermented foods (Guinee, 2004). Dairy cheeses exhibit a wide spectrum of salt content, typically ranging from 1 to 3%, with some varieties like blue cheeses exceeding 3% (Guinee & Fox, 2017). The salt content in plant-based cheeses collected in this study showed relatively low levels of salt (1.55±0.50%). This observation is likely associated with the growing focus on creating healthier products with reduced sodium content. Additionally, a label nutritional facts analysis of 245 plant-based cheese alternatives available in the United States highlighted cashew-based cheeses as having the lowest levels of sodium (Craig et al., 2022), likely due to their rich taste, potentially reducing the need for extra salt. However, it’s important to consider that consumer preferences, especially among health-conscious eaters, might also influence this trend.

In addition to the previously mentioned macronutrients and salt described on the labels, we measured the ash content in plant-based cheeses, with recorded values ranging from 2.5 to 4% (**Table S2**). These values are to be compared with the range of 1 to 7% observed in dairy cheeses, including Camembert with 3.8% ash (Fox et al., 2017). The mineral composition of cheeses is influenced by factors such as the type of milk used, the animal’s diet and the specific processing techniques. Dairy milk primarily contains calcium as the main mineral, followed by phosphorus, chloride, sodium, potassium, magnesium, and trace elements (Gaucheron, 2013). Plant-based substrates may have a similar ash percentage to dairy products, but their mineral composition can be distinct. For instance, cashew nuts contain minerals such as iron, zinc, copper, calcium, phosphorus, and magnesium (Griffin & Dean, 2017; Rico et al., 2015). Unlike dairy cheeses, the substantial presence of iron in plant-based cheese substrates can have a positive influence on microbial growth and the fermentation process.

### 3.2 Fatty acid profiles

The significant difference in fatty acid profile previously observed on the labels led us to further analyze this fact. To obtain a more detailed picture of the lipids found in plant-based cheeses collected in this study, we carried out an in-depth analysis to identify their corresponding fatty acid profiles. Our analysis revealed a predominance of oleic acid (55%) and linoleic acids (20%), which are mono-and polyunsaturated fatty acids, respectively (**Table 1**). Consistent with the label, the saturated fatty acids in our analysis represent approximately a quarter of total lipids, with on average 15% of palmitic acids and 8% of stearic acids. Tabanelli et al. (2018) analyzed raw cashew nuts and a fermented cashew food product that was spontaneously fermented and combined with salt, olive oil and lemon juice. The fatty acid profiles observed in both products of their study were similar to our findings: oleic acid constituted the predominant fat, ranging between 58 and 61%, linoleic acid ranged from 18 to 20%, palmitic acid around 10%, and stearic acid at 8%. In contrast, conventional dairy cheeses typically present around 50% of saturated long-chain fatty acid content, with a substantial proportion comprising palmitic acid, myristic acid and stearic acid (Adamska et al., 2017; Paszczyk & Łuczyńska, 2020). Additionally, dairy cheeses exhibit a diverse range of short-chain fatty acids (∼16%), including butyric acid, lauric acid, hexanoic acid, and decanoic acid. These saturated fatty acids have been associated with adverse health effects, including an increased risk of cardiovascular diseases and elevated cholesterol levels when consumed in excess (Zong et al., 2016). Although oleic acid, the main unsaturated fatty acid in plant-based cheeses, is also present in dairy cheeses, its concentration is significantly lower, reaching less than half of the amounts observed in the plant samples collected in this study (Adamska et al., 2017; Paszczyk & Łuczyńska, 2020). Oleic acid, a monounsaturated fatty acid, is known for its potential health benefits, including reducing the risk of heart disease and inflammation (Sales-Campos et al., 2013). Therefore, the higher concentration of oleic acid in plant-based cheeses may contribute to healthfulness compared to conventional dairy cheeses.

**Table 1.**
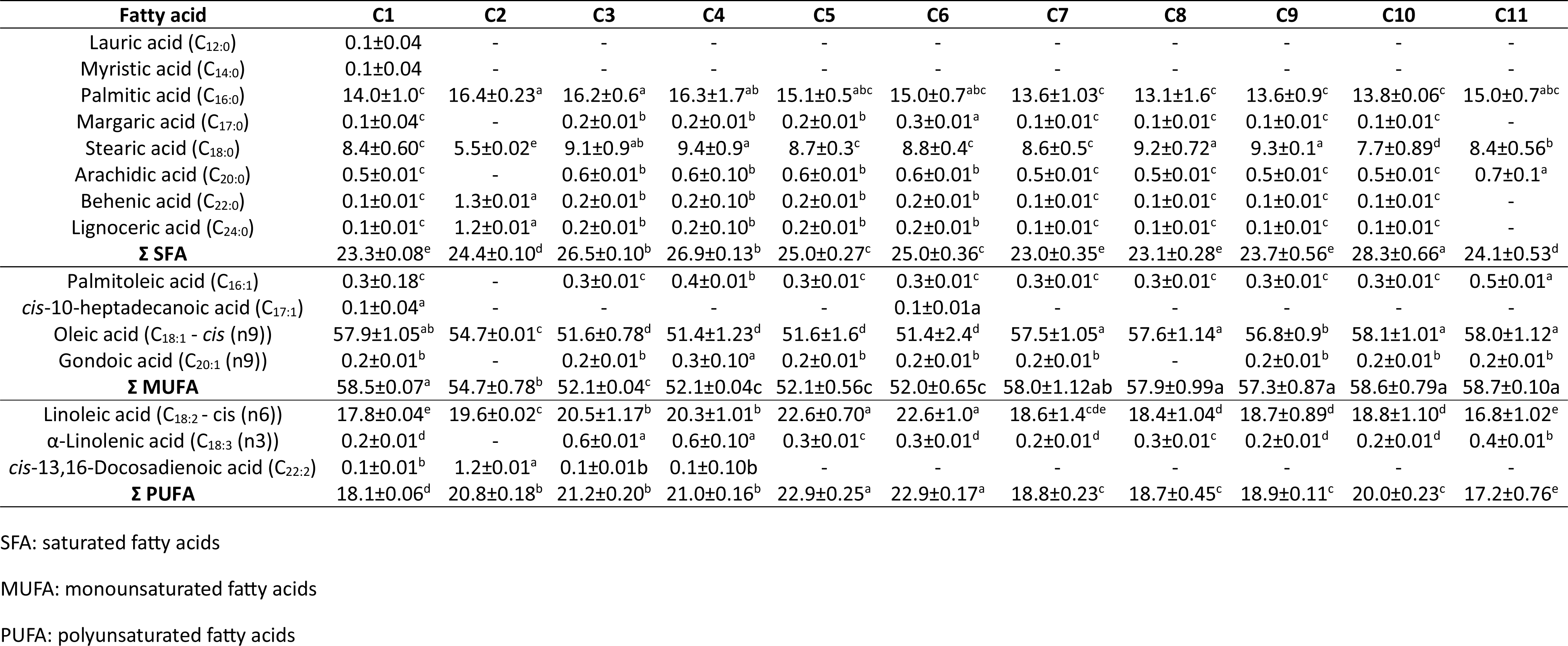
Fatty acids profiles of the 11 ripened plant-based cheese analogs collected in the European market.

### 3.3 Microbiological results

The microbial composition of fermented products is of tremendous importance in terms of safety and quality. In the following sections, we explore the microorganisms present in plant-based cheeses by culture-dependent and -independent methods to highlight their properties compared to their dairy counterparts.

#### 3.3.1 Culture-dependent composition

Overall, bacterial counts in BHI, M17, and MRS media were similar in the 11 analyzed samples (core and rind), ranging from 7.5 to 9.4 log CFU/g (**Fig. 2A and 2B**). These counts are consistent with those found in plant-based fermented foods (Ilango & Antony, 2021), but also in dairy-fermented products (da Silva Duarte et al., 2020; Kothe et al., 2021). The highest count observed in our study was in sample 8, which presents a visually distinct greenish rind. Regarding fungi present in the rinds plated on YPED, the counts ranged from 3.5 to 8.5 log CFU/g (**Fig. 2B**). This observation falls in the range found by Leclercq-Perlat et al. (2004), who explored microbiological changes in camembert-type dairy cheese by introducing various fungi, including *Penicillium camemberti* and *Geotrichum candidum*, into the model. Their study revealed that, depending on the species, the viable cell counts range from around 10^4^ CFU/g for *P. camemberti* to 10^7^ CFU/g for *G. candidum*. Interestingly, our study observed comparatively lower fungal counts in samples 5 and 6 (produced in the Netherlands), where the labels indicated the use of *P. camemberti* as a starter culture (**Table S1**).

**Fig. 2.**
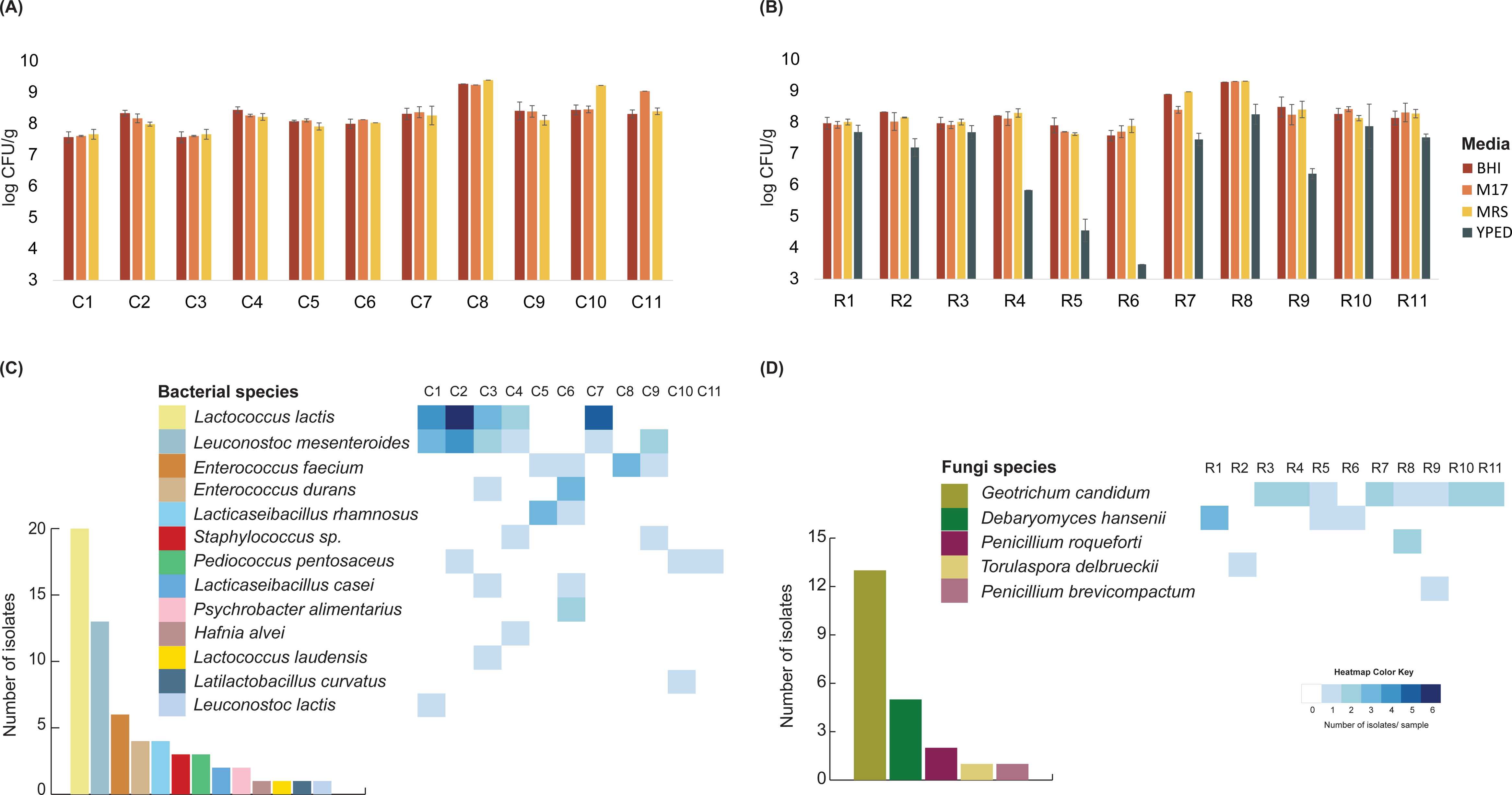
Culture-dependent results. Counts of bacterial isolates in the core (A) and rind (B) of ripened plant-based cheese analogs collected in the market, with a plot showing the number of isolates per species and a heatmap illustrating the distribution of isolates per sample for bacteria (C) and fungi (D). Samples indicated by the letter ‘C’ represent core and ‘R’ rind.

In total, we characterized 83 isolates from plant-based cheese analogs, comprising 61 bacterial and 22 fungal strains (**Table S3**). Among these isolates, we identified a variety of 15 bacterial species across the 11 samples collected. *Lactococcus lactis* constituted the predominant species (**Fig. 2C**). This prevalence recalls the dominant status of this species in the production of soft dairy cheeses (Irlinger et al., 2015; Nam et al., 2021). *L. lactis* is a species that is found in different environments, including plants, and several studies have shown the potential application of non-dairy *L. lactis* in cheese production (Cavanagh et al., 2014; McAuliffe, 2018). Future investigations including comparative genome analysis, as previously done with strains isolated in traditional fermented milk (Kothe et al., 2022), with the strains isolated from cheese analogs will be valuable in providing more information about the potential origin of these strains. Moreover, such analyses will facilitate a deeper understanding of the metabolic functions of strains tailored for plant-based alternatives, contributing to the ongoing evolution of those products.

Our study also revealed a substantial presence of *Leuconostoc mesenteroides* isolates within the plant-based cheese analogs. This species is traditionally associated with vegetable matrices (Lu et al., 2010), although in dairy cheese production, it is frequently combined with *L. lactis* (Coelho et al., 2022; Irlinger et al., 2015). The coexistence of these two species in half of the samples studied suggests a complementary metabolic relationship or functional role within the microbial community of plant-based cheeses. In fact, previous studies have already shown how beneficial the association of these species can be in terms of acidifying capacity and suitability for producing desirable aromatic compounds in the development of plant-based products (Engels et al., 2022; Harper et al., 2022). Further investigation into this relationship could provide more insights to optimize production processes for plant-based cheeses.

Furthermore, our culture-dependent approach identified several other bacterial species that could be part of microbial diversity of the plant substrates, such as *Enterococcus* spp., *Pediococcus pentosaceus, Lacticaseibacillus rhamnosus, Lacticaseibacillus casei, Latilactobacillus curvatus, and Leuconostoc lactis*. Additionally, other species commonly associated with dairy products such as *Staphylococcus*spp., *Psychrobacter alimentarius*, and *Hafnia alvei*, were isolated in this study.

Regarding fungal isolates, we identified five different species. *Geotrichum candidum*, a common starter culture in dairy bloomy cheese rinds, is predominant in most of our samples (**Fig. 2D**). Additionally, *Debaryomyces hansenii*, known for its role in dairy cheese ripening and flavor development, was isolated in three samples. Moreover, *P. roqueforti*, typically associated with dairy blue cheeses, was detected in R8 sample, where “blue” technology was utilized. The presence of *Torulaspora delbrueckii* and *Penicillium brevicompactum* in our study suggests their likely origin from the production environment and may contribute to the microbial diversity of plant-based cheese analogs, potentially influencing their quality.

Overall, the coexistence of typical dairy-associated organisms with distinct plant or environmental associated species highlights the microbial dynamics inherent to plant-based cheese production. These isolates offer the potential to improve our understanding of the microbial communities that inhabit plant-based fermented products. Their possible applications and functional properties, such as metabolic capacities, enzymatic activities and impacts on food quality and safety, could be further explored to tailor the sensory and nutritional characteristics of plant-based cheeses.

#### 3.3.2 Culture-independent composition

In this section, we further explore the taxonomic composition of plant-based cheeses collected in the European market by a culture-independent method. The bacterial and fungal composition of the 11 plant-based cheeses collected in this study was analyzed using DNA amplicon sequencing, targeting 16S and ITS genes, respectively. Clustering of the sequences resulted in 115 bacterial OTUs and 12 fungal OTUs, with taxonomic assignment up to the species level in most cases (**Table S4**). Analysis of rarefaction curves for all samples indicated saturation, suggesting sufficient sequencing depth recovery (**Fig. S1**).

Concerning bacterial composition, beta-diversity analysis revealed clustering of samples into four distinct groups (**Fig. 3A**). The first group comprised core and rind samples of plant-based cheeses produced in France with cashew nuts (C1, C2, R2, C4, R4, and C7), with the most abundant species being *L. lactis* (**Fig. 3B**). The second group comprised samples 8 and 9 from the same producer in France and presented a predominance of *L. mesenteroides*. The third group included samples from the Netherlands (samples 5 and 6), predominantly containing *Lactobacillus acidophilus, Bifidobacterium animalis, Enterococcus sp.,* and *Streptococcus thermophilus.* The last group is not homogenous and consisted of a sample made with almond milk (C10, R10), along with rinds from R1, R3 and R7 samples, exhibiting diverse species composition. Sample 10 showed similar compositions in its core and rind, dominated by *Latilactobacillus sakei, L. lactis, Leuconostoc spp., and Pediococcus spp*., while the rinds of the other samples in this group contained *Enterobacteriaceae* and potential spoilage species such as *Qipengyuania sp.* and *Pseudomonas sp*.

**Fig. 3.**
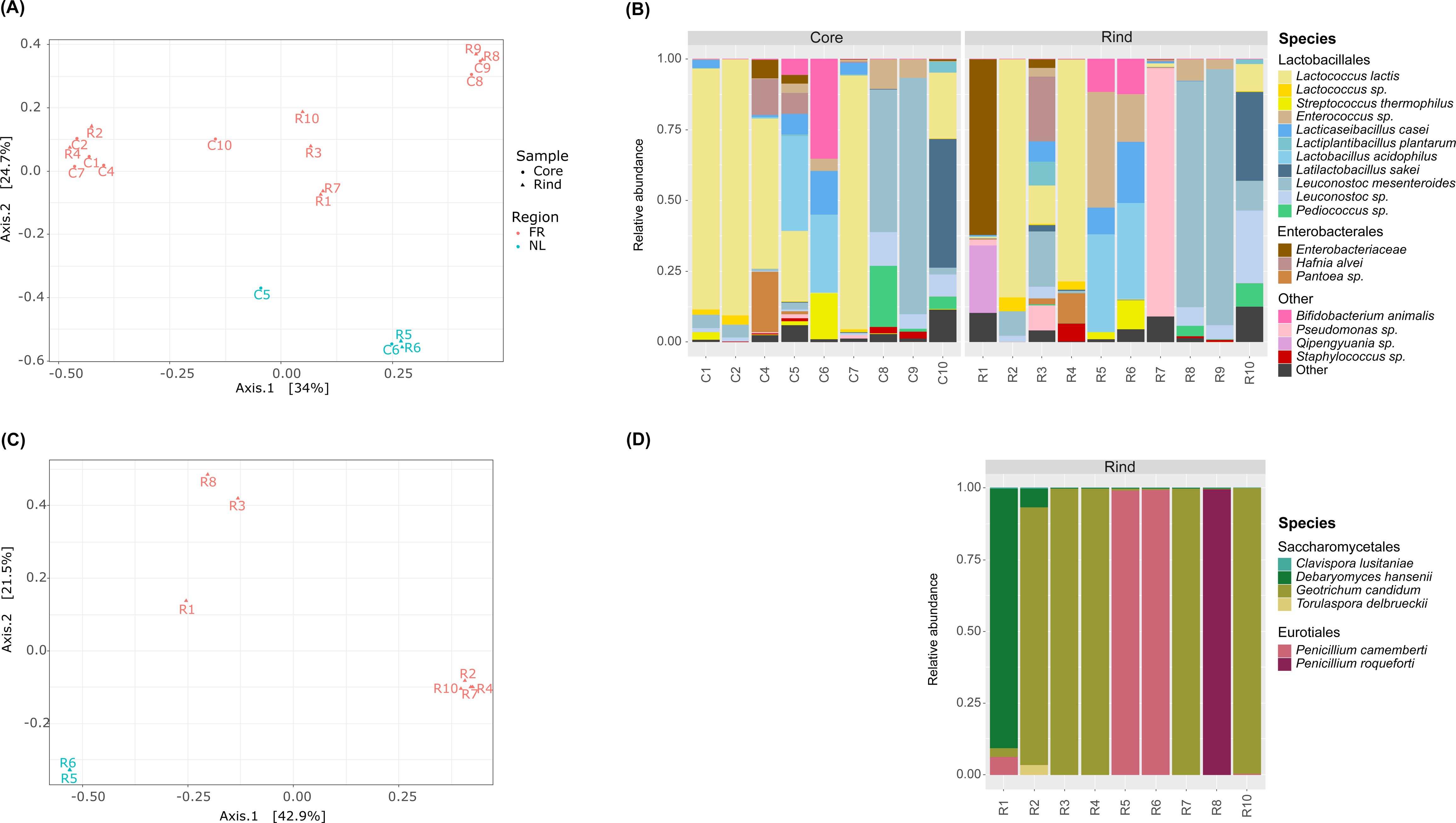
Principal Coordinate Analysis (PCoAs) and composition plots illustrating the bacterial communities (A, B) and fungal communities (C, D) identified in the ripened plant-based cheese analogs.

Several species found in plant-based cheese are similar to those of starter cultures used in dairy products, including potential probiotics and bacteria contributing to aroma and flavor formation. The dominance of several of these species suggests they were added as starters in the plant-based cheese, while less abundant species may originate from plant matrices or the production environment. For instance, *L. lactis,* identified as a primary bacterium in many starter cultures for dairy cheeses, was abundant in plant cheeses collected from different producers in France. While traditionally associated with dairy (Mills et al., 2011), *L. lactis* also occurs, as mentioned previously, in non-dairy niches like plant-based fermented foods (Gustaw et al., 2021). Similarly, *L. mesenteroides*, another common lactic acid bacterium in food fermentation, was found in significant amounts in certain French cheeses. While primarily used in dairy products, some strains of *L. mesenteroides* also possess probiotic potential and are used in various fermented foods, mainly those using vegetables like sauerkraut and kimchi (Chun et al., 2017).

In sample C10, a different taxonomic composition was observed, with co-dominance of *Lactobacillus sakei* and *L. lactis*. This sample also exhibited a unique cylindrical shape similar to traditional “bouchon de chèvre” cheese (**Fig. 1A**), which is a surface-mold cheese produced from goat’s milk. Although often studied in meat and seafood products (Chaillou et al., 2015; Zagorec & Champomier-Vergès, 2017), *L. sakei* has been found in plant-based fermented foods (Landis et al., 2021; Xiao et al., 2020), suggesting its adaptability across different niches.

Samples 5 and 6 produced in the Netherlands displayed a specific composition, with the dominance of *L. acidophilus* and *B. animalis*, which are typical probiotic strains also used in plant-based milk substitutes (Rasika et al., 2021). These strains, not naturally present in plants, are likely industrial starters. Additionally, dominance of *Enterococcus* spp. on the plant cheese rind suggests its presence could be attributed to the industrial environment or human origin (Ben Said et al., 2016; Krawczyk et al., 2021; Terzić-Vidojević et al., 2020).

Regarding the fungal composition, beta-diversity analysis showed clustering of rinds into three distinct groups (**Fig. 3C**). The first group comprised samples from the Netherlands, with *P. camemberti* being the most abundant species (**Fig. 3D**). The second group was predominantly containing *G. candidum* (distinct OTU from the R3 sample, **Table S4**). The remaining rind samples included R1, R3, and R8, each exhibiting a diverse species composition. For instance, R1 is rich in *D. hansenii*, R3 in *G. candidum*, and R8 is predominantly in *P. roqueforti*. All these species are among the most frequent yeast/fungi starters utilized in the dairy industry. *Geotrichum candidum* and *P. camemberti* are domesticated species commonly used in the production of soft cheeses with bloomy rinds, and they contribute to cheese flavor (Ropars & Giraud, 2022). *Debaryomyces hansenii*, a halophilic yeast species, is commonly found on the rinds of various cheese varieties (Wolfe et al., 2014). And *P. roqueforti* is a filamentous fungus that is widely used in various types of blue cheeses.

Overall, while several species present in plant-based cheeses may originate from plants or environmental sources, the predominant ones were likely added from industrial starters also used in dairy productions. The presence of various typical dairy cheese species in plant-based cheese products highlights their adaptability to this new niche, suggesting a purposeful effort to reproduce the intricate flavors and textures of traditional dairy cheeses. Due to their beneficial attributes, we hypothesize that these dairy cheese species were intentionally introduced into these plant-based cheeses, aimed at enhancing the sensory experience of these products.

## 4. Conclusion & Perspectives

This research contributes to expanding our understanding of the nutritional composition and microbial ecology of ripened plant-based cheese analogs present on the European market, particularly those produced from cashew nuts. Our analyses revealed a macronutrient profile rich in lipids, proteins, and carbohydrates, with notable differences in fatty acid composition compared to traditional dairy cheeses. Specifically, the predominance of oleic acid in plant-based cheeses, a monounsaturated fatty acid, diverges from the prevalence of saturated fatty acids in dairy cheeses, impacting both health considerations and sensory attributes.

While several strains identified in our study may originate from plant or environment sources, most others likely originated from dairy starters. The presence of typical dairy cheese species in plant-based ones likely highlights the adaptability of several selected strains to this emerging niche. It might also reflect the lack of proper starter strains isolated from plants that would be able to develop products with good sensorial quality from vegetal raw material.

It is important to acknowledge that the plant cheese alternatives analyzed in this study are derived from cashew nuts and almonds, substrates known for their high cost and limited sustainability in agricultural practices (Cap et al., 2023). While these ingredients offer advantages in cheese-making technologies, attributed to their neutral flavor profile and absence of undesirable tastes, broader investigation into alternative plant ingredients is important to address sustainability challenges and explore novel flavor profiles.

Addressing the differences observed compared to dairy cheeses presents a fundamental challenge in plant-based food production. Exploring fermentation processes and utilizing mixed plant-based ingredients offer promising approaches to mimic traditional dairy-based cheese features. Future research could delve into genomic analyses to understand microbial adaptations to plant matrices and shed light on ecological dynamics. By leveraging interdisciplinary approaches and innovative technologies, such as those in microbiology and chemistry, we can advance the development of plant-based cheeses that meet consumer preferences while contributing to a more sustainable food future. These findings pave the way for product development and deepen our scientific understanding of plant-based alternatives, fostering innovation and progress in this expanding field.

## Supporting information

Table S3 Bacteria

Table S3 Fungi

Table S4 Bacteria

Table S4 Fungi

## Supplementary Material

**Fig. S1.** Rarefaction curves depicting the sequencing depth of 16S (A) and ITS (B) and species richness for the core and rind samples obtained from plant-based cheeses.

**Table S1.** Metadata of the plant-based cheese analogs collected showing the geographical location of production, producer, ingredients, rind type, technology and photo.

**Table S2.** Nutritional composition of the ripened plant-based cheeses according to nutritional facts on the labels and protein and ash measurements.

**Table S3.** Bacterial and fungal isolates from 11 ripened plant-based cheese analogs collected in the market, including their respective growth medium, sequence, and best blast hit for characterization. The excel file is divided into two sheets: ‘Bacteria’ and ‘Fungi’.

**Table S4.** Number of raw reads detected for bacterial and fungal operational taxonomic units (OTUs) in the samples. The color code indicates the reads mapped for each species, ranging from red (low abundance) to green (high abundance). The excel file is divided into two sheets: ‘16S’ and ‘ITS’.

## Acknowledgements

We would like to thank Dra. Michele Bertoni Mann and Dra. Ana Murtele for the technical support in sequencing, as well Andressa Duprat for the technical support in the fatty acid profiles. CIK and JE have been funded by The Novo Nordisk Foundation, grant number NNF20CC0035580. JF has been funded by CNPq #307810/2021-6.

## Author Contributions

CIK designed the experiment. IJ performed microbiology related experiments. CRK performed the fatty acid analyses. IJ and CIK wrote the first draft of the manuscript. CIK performed microbial analyses and graphs. CIK and PR elaborated the points for discussion. JF, PR, JDE and CIK revised the manuscript. PR, JDE and JF provided financial support for the research. All authors had the opportunity to read, comment on, and approve the manuscript before submission.

