## Supplementary material for "Ripened plant-based cheese analogs in Europe: nutritional and microbial profiles": Table S2

Table of nutritional facts on plant-based cheese analogs based on labels

| | **C1** | **C2** | **C3** | **C4** | **C5** | **C6** | **C7** | **C8** | **C9** | **C10** | **C11** |
|---|---|---|---|---|---|---|---|---|---|---|---|
| **Energy (Kcal)** | 379 | 377 | 214 | 379 | 376 | 376 | 375 | 528.9 | 492.9 | 389 | 286 |
| **Lipid (g)** | 30 | 29 | 17 | 31 | 31 | 31.3 | 29.9 | 41.5 | 38.23 | 33 | 20 |
| **Saturated fatty acids (g)** | 7.3 | 6.4 | 4.5 | 6.4 | 6.3 | 6.3 | 4.8 | 11.7 | 8.17 | 6.9 | 4 |
| **Carbohydrates (g)** | 15 | 15 | 5.5 | 10.3 | 10.3 | 10.3 | 13.6 | 16.8 | 16.02 | 7.3 | 12 |
| ***of which no sugar*** | <0.5 | <0.5 | <0.5 | <0.5 | <0.5 | <0.5 | 2.7 | 6.9 | 6.2 | - | 3 |
| **Protein (g)** | 14 | 14 | 10 | 13.6 | 13.7 | 13.7 | 13.2 | 15.2 | 14.6 | 14 | 10 |
| **Salt (g)** | 1.2 | 0.77 | 1.2 | 1.75 | 1.77 | 1.77 | 0.8 | 2.2 | 2.1 | 1.5 | 2 |

Protein and ash measures on plant-based cheese analogs

| | **C1** | **C2** | **C3** | **C4** | **C5** | **C6** | **C7** | **C8** | **C9** | **C10** | **C11** |
|---|---|---|---|---|---|---|---|---|---|---|---|
| Ash (%)* | 3.12 | 3.54 | 2.48 | 3.07 | 2.86 | 3.99 | 2.44 | 3.23 | 4.17 | 2.97 | 3.02 |
| Protein (%)* | 13.22 | 11.47 | 9.04 | 9.68 | 10.66 | 12.04 | 13.84 | 12.73 | 13.51 | 12.01 | 13.15 |

*Average of duplicate

| Average | Standard deviation |
|---|---|
| 379.35 | 84.53 |
| 30.18 | 6.95 |
| 6.62 | 2.09 |
| 12.01 | 3.64 |
| 4.70 | 2.16 |
| 13.27 | 1.70 |
| 1.55 | 0.50 |

| Average | Standard deviation |
|---|---|
| 3.171818 | 0.54708 |
| 11.94091 | 1.587239 |
