## Supplementary material for "Ripened plant-based cheese analogs in Europe: nutritional and microbial profiles": Table S3 Bacteria

| Sample | Part | Media | Clone | Species, Best hit | Per. Ident. (%) | Query Length |
|---|---|---|---|---|---|---|
| C1 | Core | BHI | C1C1 | *Lactococcus lactis* | 99.83 | 1207 |
| C1 | Core | BHI | C1C2 | *Lactococcus lactis* | 99.84 | 1214 |
| C1 | Core | BHI | C1C4 | *Leuconostoc lactis* | 100.00 | 1156 |
| C1 | Core | M17 | C1C5 | *Leuconostoc mesenteroides* | 100.00 | 1156 |
| C1 | Core | M17 | C1C7 | *Lactococcus lactis* | 99.92 | 2370 |
| C1 | Core | MRS | C1C11 | *Leuconostoc mesenteroides* | 99.74 | 760 |
| C1 | Core | MRS | C1C12 | *Lactococcus lactis* | 99.92 | 1186 |
| C1 | Rind | MRS | C1R16 | *Leuconostoc mesenteroides* | 99.83 | 1209 |
| C2 | Core | BHI | C2C2 | *Lactococcus lactis* | 99.59 | 2408 |
| C2 | Core | BHI | C2C3 | *Leuconostoc mesenteroides* | 99.65 | 1157 |
| C2 | Core | BHI | C2C4 | *Leuconostoc mesenteroides* | 99.91 | 1153 |
| C2 | Core | M17 | C2C7 | *Lactococcus lactis* | 99.67 | 1217 |
| C2 | Core | M17 | C2C8 | *Leuconostoc mesenteroides* | 99.83 | 1189 |
| C2 | Core | MRS | C2C11 | *Pediococcus pentosaceus* | 99.75 | 1214 |
| C2 | Core | MRS | C2C12 | *Lactococcus lactis* | 99.91 | 1175 |
| C2 | Rind | M17 | C2R3 | *Leuconostoc mesenteroides* | 99.85 | 676 |
| C2 | Rind | M17 | C2R4 | *Lactococcus lactis* | 99.75 | 1207 |
| C2 | Rind | BHI | C2R9 | *Lactococcus laudensis* | 99.79 | 962 |
| C2 | Rind | BHI | C2R11 | *Lactococcus lactis* | 99.92 | 1203 |
| C2 | Rind | MRS | C2R15 | *Lactococcus lactis* | 99.75 | 1208 |
| C3 | Core | MRS | C3C10 | *Leuconostoc mesenteroides* | 100.00 | 1154 |
| C3 | Rind | BHI | C3C1 | *Lactococcus lactis* | 99.77 | 862 |
| C3 | Rind | BHI | C3C4 | *Lactococcus lactis* | 99.72 | 1064 |
| C3 | Rind | M17 | C3C5 | *Lactococcus lactis* | 99.82 | 1107 |
| C3 | Rind | M17 | C3C7 | *Leuconostoc mesenteroides* | 99.81 | 1066 |
| C3 | Rind | MRS | C3C12 | *Lacticaseibacillus paracasei* | 99.65 | 857 |
| C3 | Rind | M17 | C4R8 | *Enterococcus durans* | 99.64 | 554 |
| C4 | Core | MRS | C4R11 | *Leuconostoc mesenteroides* | 99.91 | 1153 |
| C4 | Core | BHI | C4R13 | *Staphylococcus succinus* | 99.58 | 1184 |
| C4 | Core | BHI | C4R14 | *Lactococcus lactis* | 99.83 | 1190 |
| C4 | Rind | M17 | C3R8 | *Lactococcus lactis* | 98.96 | 671 |
| C4 | Rind | M17 | C4C8 | *Hafnia alvei* | 99.72 | 361 |
| C5 | Core | M17 | C5C9 | *Lacticaseibacillus rhamnosus* | 99.04 | 1045 |
| C5 | Core | MRS | C5C8 | *Lacticaseibacillus rhamnosus* | 99.39 | 825 |
| C5 | Core | BHI | C5C3 | *Lacticaseibacillus rhamnosus* | 99.42 | 1197 |
| C5 | Rind | BHI | C5R1 | *Enterococcus faecium* | 99.32 | 886 |
| C6 | Core | M17 | C6C9 | *Enterococcus faecium* | 99.74 | 1173 |
| C6 | Core | M17 | C6C12 | *Lacticaseibacillus rhamnosus* | 100.00 | 1178 |
| C6 | Core | BHI | C6C2 | *Enterococcus durans* | 99.72 | 1060 |
| C6 | Rind | MRS | C6R7 | *Enterococcus durans* | 99.81 | 1066 |
| C6 | Rind | MRS | C6R5 | *Psychrobacter alimentarius* | 99.63 | 1079 |
| C6 | Rind | BHI | C6R1 | *Psychrobacter alimentarius* | 99.45 | 1097 |
| C6 | Rind | BHI | C6R2 | *Enterococcus durans* | 99.63 | 1075 |
| C7 | Core | M17 | C7C9 | *Lactococcus lactis* | 99.75 | 1186 |
| C7 | Core | M17 | C7C12 | *Lactococcus lactis* | 99.83 | 1176 |
| C7 | Core | MRS | C7C5 | *Lactococcus lactis* | 99.75 | 1202 |
| C7 | Core | BHI | C7C4 | *Lactococcus lactis* | 99.30 | 869 |
| C7 | Rind | MRS | C7R7 | *Lacticaseibacillus casei* | 99.73 | 746 |
| C7 | Rind | M17 | C7R10 | *Leuconostoc mesenteroides* | 99.72 | 721 |
| C7 | Rind | MRS | C7R6 | *Lactococcus lactis* | 99.70 | 674 |
| C8 | Core | M17 | C8-C4 | *Enterococcus faecium* | 99.91 | 1171 |
| C8 | Core | M17 | C8-C10 | *Enterococcus faecium* | 99.33 | 1193 |
| C8 | Rind | M17 | C8-R2 | *Enterococcus faecium* | 99.40 | 1164 |
| C9 | Core | M17 | C9-C1 | *Enterococcus faecium* | 99.66 | 1173 |
| C9 | Core | M17 | C9-C5 | *Leuconostoc mesenteroides* | 99.91 | 1159 |
| C9 | Rind | M17 | C9-R1 | *Leuconostoc mesenteroides* | 99.74 | 1148 |
| C9 | Rind | M17 | C9-R4 | *Staphylococcus saprophyticus* | 99.00 | 1202 |
| C9 | Rind | M17 | C9-R9 | *Staphylococcus sp.* | 95.57 | 1147 |
| C10 | Rind | M17 | C10-R2 | *Pediococcus pentosaceus* | 98.90 | 1183 |
| C10 | Rind | M17 | C10-R11 | *Latilactobacillus curvatus* | 98.74 | 1192 |
| C11 | Core | M17 | C11-C11 | *Pediococcus pentosaceus* | 99.23 | 1173 |
