## Supplementary material for "Ripened plant-based cheese analogs in Europe: nutritional and microbial profiles": Table S3 Fungi

| Sample | Part | Media | Clone | Species | Per. Ident. (%) | Query Length |
|---|---|---|---|---|---|---|
| C1 | Rind | YPED | C1R3 | *Debaryomyces hansenii* | 100.00 | 578 |
| C1 | Rind | YPED | C1R5 | *Debaryomyces hansenii* | 100.00 | 576 |
| C1 | Rind | YPED | C1R6 | *Debaryomyces hansenii* | 99.66 | 581 |
| C2 | Rind | YPED | C2R14 | *Torulaspora delbrueckii* | 99.32 | 594 |
| C3 | Rind | YPED | R3 10-3 | *Geotrichum candidum* | 97.58 | 254 |
| C3 | Rind | YPED | R3 10-5 | *Geotrichum candidum* | 99.38 | 555 |
| C4 | Rind | YPED | R4(1) | *Geotrichum candidum* | 98.90 | 547 |
| C4 | Rind | YPED | R4(2) | *Geotrichum candidum* | 98.92 | 552 |
| C5 | Rind | YPED | C5R16 | *Geotrichum candidum* | 99.29 | 560 |
| C5 | Rind | YPED | C5R13 | *Debaryomyces hansenii* | 100.00 | 577 |
| C6 | Rind | YPED | C6R14 | *Debaryomyces hansenii* | 99.66 | 579 |
| C7 | Rind | YPED | C7R12 | *Geotrichum candidum* | 99.63 | 545 |
| C7 | Rind | YPED | C7R5 | *Geotrichum candidum* | 98.71 | 232 |
| C8 | Rind | YPED | C8R16 | *Geotrichum candidum* | 99.64 | 560 |
| C8 | Rind | YPED | C8R15 | *Penicillium roqueforti* | 99.30 | 576 |
| C8 | Rind | YPED | C8R12 | *Penicillium roqueforti* | 98.80 | 337 |
| C9 | Rind | YPED | C9R14 | *Geotrichum candidum* | 99.29 | 564 |
| C9 | Rind | YPED | C9R15 | *Penicillium brevicompactum* | 100.00 | 388 |
| C10 | Rind | YPED | C10R12 | *Geotrichum candidum* | 99.27 | 546 |
| C10 | Rind | YPED | C10-R11 | *Geotrichum candidum* | 98.69 | 539 |
| C11 | Rind | YPED | C11R2 | *Geotrichum candidum* | 99.81 | 546 |
| C11 | Rind | YPED | C11R12 | *Geotrichum candidum* | 99.08 | 542 |
