## Supplementary material for "Ripened plant-based cheese analogs in Europe: nutritional and microbial profiles": Table S4 Bacteria

| Kingdom | Phylum | Class | Oder | Family | Genus | Species | seed_sequ | observation_ | observation | C1 | C2 | C4 | C5 | C6 | C7 | C8 | C9 | C10 | R1 | R2 | R3 | R4 | R5 | R6 | R7 | R8 | R9 | R10 |
|---|---|---|---|---|---|---|---|---|---|---|---|---|---|---|---|---|---|---|---|---|---|---|---|---|---|---|---|---|
| Bacteria | Firmicutes | Bacilli | Lactobacillales | Streptococcaceae | Lactococcus | Lactococcus lactis | TACGTAG | Cluster_1 | 285045 | 28129 | 24773 | 30610 | 9799 | 26 | 62322 | 10 | 10 | 15757 | 499 | 35374 | 7086 | 68192 | 15 | 46 | 788 | 15 | 4 | 1590 |
| Bacteria | Firmicutes | Bacilli | Lactobacillales | Lactobacillaceae | Leuconostoc | Leuconostoc mesenteroides | TACGTAT | Cluster_2 | 186962 | 1416 | 1233 | 518 | 919 | 0 | 236 | 29591 | 24714 | 1677 | 108 | 3326 | 9777 | 512 | 2 | 0 | 56 | 53786 | 57372 | 1719 |
| Bacteria | Proteobac | Gammaproteo | Enterobacterales | Enterobacteriaceae | Escherichia-Shigella | Enterobacteriaceae | TACGGAG | Cluster_3 | 64679 | 0 | 1 | 3638 | 1178 | 1 | 62 | 2 | 0 | 409 | 57693 | 0 | 1583 | 4 | 0 | 0 | 102 | 4 | 2 | 0 |
| Bacteria | Proteobac | Gammaproteo | Pseudomonadale | Pseudomonadaceae | Pseudomonas | Pseudomonas sp. | TACAGAG | Cluster_4 | 49298 | 0 | 0 | 20 | 150 | 0 | 248 | 10 | 9 | 96 | 10 | 3 | 37 | 9 | 0 | 0 | 48696 | 6 | 4 | 0 |
| Bacteria | Firmicutes | Bacilli | Lactobacillales | Lactobacillaceae | Lactobacillus | Lactobacillus acidophilus | TACGTAG | Cluster_5 | 34692 | 2 | 2 | 189 | 13083 | 8547 | 8 | 27 | 10 | 12 | 180 | 4 | 6 | 7 | 6323 | 6264 | 8 | 12 | 6 | 2 |
| Bacteria | Firmicutes | Bacilli | Lactobacillales | Lactobacillaceae | Latilactobacillus | Latilactobacillus sakei | TACGTAG | Cluster_6 | 37015 | 10 | 0 | 66 | 86 | 14 | 123 | 68 | 3 | 29956 | 0 | 1 | 1212 | 312 | 6 | 10 | 6 | 6 | 4 | 5132 |
| Bacteria | Firmicutes | Bacilli | Lactobacillales | Enterococcaceae | Enterococcus | Enterococcus sp. | TACGTAG | Cluster_7 | 27406 | 15 | 1 | 186 | 1172 | 1151 | 527 | 5368 | 1712 | 39 | 0 | 2 | 1162 | 106 | 7275 | 2618 | 103 | 4537 | 1431 | 1 |
| Bacteria | Proteobac | Gammaproteo | Enterobacterales | Hafniaceae | Hafnia | Hafnia alvei | TACGGAG | Cluster_8 | 22981 | 7 | 1 | 7571 | 2915 | 0 | 161 | 2 | 0 | 55 | 1 | 1 | 11986 | 3 | 1 | 0 | 272 | 2 | 1 | 2 |
| Bacteria | Proteobac | Gammaproteo | Enterobacterales | Erwiniaceae | Pantoea | Pantoea sp. | TACGGAG | Cluster_9 | 17615 | 0 | 0 | 9229 | 302 | 0 | 4 | 2 | 0 | 3 | 11 | 1 | 1053 | 6905 | 0 | 0 | 103 | 0 | 2 | 0 |
| Bacteria | Actinobac | Actinobacteria | Bifidobacteriales | Bifidobacteriaceae | Bifidobacterium | Bifidobacterium animalis | TACGTAG | Cluster_10 | 17612 | 0 | 0 | 149 | 2156 | 10760 | 0 | 0 | 0 | 21 | 0 | 0 | 0 | 0 | 2149 | 2374 | 0 | 3 | 0 | 0 |
| Bacteria | Firmicutes | Bacilli | Lactobacillales | Lactobacillaceae | Pediococcus | Pediococcus sp. | TACGTAG | Cluster_11 | 20379 | 27 | 1 | 59 | 29 | 0 | 1 | 13043 | 309 | 2922 | 0 | 1 | 188 | 0 | 0 | 0 | 1 | 2299 | 167 | 1332 |
| Bacteria | Firmicutes | Bacilli | Lactobacillales | Lactobacillaceae | Leuconostoc | Leuconostoc sp. | TACGTAT | Cluster_12 | 13281 | 28 | 5 | 25 | 149 | 0 | 320 | 1149 | 89 | 5165 | 0 | 1 | 9 | 1 | 0 | 0 | 131 | 1906 | 277 | 4026 |
| Bacteria | Firmicutes | Bacilli | Lactobacillales | Lactobacillaceae | Lacticaseibacillus | Lacticaseibacillus casei | TACGTAG | Cluster_13 | 21065 | 910 | 0 | 311 | 2910 | 4417 | 2582 | 1 | 0 | 9 | 332 | 0 | 3615 | 12 | 1643 | 3855 | 459 | 4 | 1 | 4 |
| Bacteria | Proteobac | Gammaproteo | Pseudomonadale | Pseudomonadaceae | Pseudomonas | Pseudomonas sp. | TACAGAG | Cluster_14 | 12901 | 0 | 0 | 49 | 311 | 0 | 49 | 1 | 1 | 11 | 1177 | 1 | 4195 | 1 | 0 | 0 | 7101 | 3 | 1 | 0 |
| Bacteria | Firmicutes | Bacilli | Staphylococcales | Staphylococcaceae | Staphylococcus | Staphylococcus sp. | TACGTAG | Cluster_15 | 8770 | 17 | 58 | 292 | 405 | 0 | 2 | 1246 | 681 | 35 | 0 | 27 | 1 | 5177 | 0 | 4 | 0 | 380 | 445 | 0 |
| Bacteria | Firmicutes | Bacilli | Lactobacillales | Lactobacillaceae | Lactiplantibacillus | Lactiplantibacillus plantarum | TACGTAG | Cluster_16 | 7960 | 7 | 0 | 124 | 289 | 3 | 325 | 0 | 1 | 2455 | 0 | 0 | 4414 | 0 | 4 | 4 | 52 | 0 | 0 | 282 |
| Bacteria | Firmicutes | Bacilli | Lactobacillales | Streptococcaceae | Streptococcus | Streptococcus thermophilus | TACGTAG | Cluster_17 | 9383 | 928 | 1 | 211 | 619 | 5094 | 8 | 64 | 57 | 7 | 0 | 0 | 40 | 0 | 481 | 1870 | 1 | 2 | 0 | 0 |
| Bacteria | Proteobac | Alphaproteoba | Sphingomonadal | Sphingomonadaceae | Qipengyuania | Qipengyuania sp. | TACGGAG | Cluster_18 | 18557 | 0 | 0 | 0 | 0 | 0 | 0 | 0 | 0 | 0 | 18557 | 0 | 0 | 0 | 0 | 0 | 0 | 0 | 0 | 0 |
| Bacteria | Proteobac | Gammaproteo | Enterobacterales | Erwiniaceae | Pantoea | Pantoea sp. | TACGGAG | Cluster_19 | 6032 | 0 | 0 | 3482 | 74 | 0 | 2 | 1 | 0 | 1 | 395 | 0 | 1 | 2058 | 0 | 0 | 18 | 0 | 0 | 0 |
| Bacteria | Firmicutes | Bacilli | Lactobacillales | Lactobacillaceae | Weissella | Weissella sp. | TACGTAT | Cluster_20 | 5185 | 0 | 0 | 3 | 290 | 1 | 1 | 823 | 2 | 2231 | 268 | 0 | 0 | 145 | 0 | 0 | 1 | 597 | 0 | 823 |
| Bacteria | Firmicutes | Bacilli | Lactobacillales | Lactobacillaceae | Weissella | Weissella viridescens | TACGTAT | Cluster_21 | 4528 | 0 | 0 | 0 | 16 | 0 | 0 | 2 | 0 | 4217 | 0 | 0 | 0 | 0 | 0 | 0 | 0 | 1 | 0 | 292 |
| Bacteria | Proteobac | Gammaproteo | Enterobacterales | Yersiniaceae | Yersinia | Yersinia sp. | TACGGAG | Cluster_22 | 3936 | 0 | 0 | 0 | 19 | 0 | 332 | 0 | 1 | 2 | 1 | 1 | 1 | 2 | 0 | 0 | 3575 | 1 | 1 | 0 |
| Bacteria | Firmicutes | Bacilli | Lactobacillales | Lactobacillaceae | Leuconostoc | Leuconostoc sp. | TACGTAG | Cluster_23 | 6698 | 62 | 0 | 0 | 1 | 0 | 4 | 2505 | 852 | 35 | 0 | 1 | 560 | 7 | 0 | 0 | 0 | 1250 | 1426 | 55 |
| Bacteria | Proteobac | Alphaproteoba | Sphingomonadal | Sphingomonadaceae | Qipengyuania | Qipengyuania sp. | TACGGAG | Cluster_24 | 3434 | 0 | 0 | 0 | 0 | 0 | 0 | 0 | 0 | 0 | 3434 | 0 | 0 | 0 | 0 | 0 | 0 | 0 | 0 | 0 |
| Bacteria | Proteobac | Gammaproteo | Enterobacterales | Yersiniaceae | Serratia | Serratia sp. | TACGGAG | Cluster_25 | 2340 | 0 | 0 | 19 | 17 | 1 | 17 | 0 | 0 | 2 | 0 | 0 | 0 | 0 | 0 | 0 | 2281 | 0 | 0 | 3 |
| Bacteria | Firmicutes | Bacilli | Lactobacillales | Lactobacillaceae | Leuconostoc | Leuconostoc sp. | TACGTAT | Cluster_26 | 4029 | 70 | 0 | 0 | 1 | 0 | 11 | 2219 | 114 | 36 | 0 | 0 | 770 | 0 | 0 | 0 | 1 | 487 | 216 | 104 |
| Bacteria | Proteobac | Gammaproteo | Pseudomonadale | Pseudomonadaceae | Pseudomonas | Pseudomonas sp. | TACAGAG | Cluster_27 | 2888 | 0 | 0 | 45 | 41 | 0 | 92 | 0 | 0 | 0 | 470 | 2 | 319 | 0 | 0 | 0 | 1919 | 0 | 0 | 0 |
| Bacteria | Firmicutes | Bacilli | Lactobacillales | Lactobacillaceae | Leuconostoc | Leuconostoc mesenteroides | TACGTAT | Cluster_28 | 3687 | 129 | 68 | 0 | 1 | 0 | 15 | 1377 | 340 | 12 | 0 | 201 | 167 | 44 | 0 | 0 | 0 | 669 | 647 | 17 |
| Bacteria | Proteobac | Alphaproteoba | Sphingomonadal | Sphingomonadaceae | Sphingomicrobium | Sphingomicrobium sp. | TACGGAG | Cluster_29 | 4290 | 0 | 0 | 0 | 0 | 0 | 0 | 0 | 0 | 0 | 4290 | 0 | 0 | 0 | 0 | 0 | 0 | 0 | 0 | 0 |
| Bacteria | Firmicutes | Bacilli | Lactobacillales | Lactobacillaceae | Leuconostoc | Leuconostoc sp. | TACGTAT | Cluster_30 | 4174 | 46 | 22 | 0 | 1 | 0 | 3 | 1280 | 550 | 3 | 0 | 58 | 141 | 8 | 1 | 0 | 1 | 967 | 1086 | 7 |
| Bacteria | Proteobac | Gammaproteo | Pseudomonadale | Moraxellaceae | Acinetobacter | Acinetobacter johnsonii | TACAGAG | Cluster_31 | 1540 | 0 | 0 | 309 | 7 | 0 | 31 | 1 | 0 | 25 | 1017 | 0 | 149 | 0 | 0 | 0 | 1 | 0 | 0 | 0 |
| Bacteria | Firmicutes | Bacilli | Lactobacillales | Enterococcaceae | Enterococcus | Enterococcus sp. | TACGTAT | Cluster_32 | 1326 | 0 | 0 | 0 | 1 | 0 | 3 | 387 | 200 | 0 | 0 | 0 | 37 | 0 | 0 | 1 | 0 | 393 | 304 | 0 |
| Bacteria | Firmicutes | Bacilli | Lactobacillales | Lactobacillaceae | Leuconostoc | Leuconostoc sp. | TACGTAG | Cluster_33 | 1921 | 242 | 345 | 0 | 13 | 1 | 29 | 10 | 7 | 29 | 0 | 872 | 266 | 61 | 0 | 0 | 1 | 7 | 2 | 36 |
| Bacteria | Firmicutes | Bacilli | Lactobacillales | Streptococcaceae | Lactococcus | Lactococcus sp. | TACGTAG | Cluster_34 | 2053 | 0 | 696 | 0 | 100 | 0 | 0 | 1 | 1 | 0 | 0 | 1255 | 0 | 0 | 0 | 0 | 0 | 0 | 0 | 0 |
| Bacteria | Firmicutes | Bacilli | Lactobacillales | Streptococcaceae | Lactococcus | Lactococcus sp. | TACGTAG | Cluster_35 | 2050 | 14 | 59 | 0 | 1 | 0 | 1 | 0 | 0 | 3 | 0 | 28 | 0 | 1944 | 0 | 0 | 0 | 0 | 0 | 0 |
| Bacteria | Firmicutes | Bacilli | Bacillales | Planococcaceae | Kurthia | Kurthia sp. | TACGTAG | Cluster_36 | 1053 | 0 | 0 | 0 | 43 | 0 | 0 | 0 | 0 | 0 | 0 | 1 | 1009 | 0 | 0 | 0 | 0 | 0 | 0 | 0 |
| Bacteria | Firmicutes | Bacilli | Lactobacillales | Enterococcaceae | Enterococcus | Enterococcus sp. | TACGTAG | Cluster_37 | 971 | 0 | 0 | 0 | 0 | 99 | 7 | 313 | 3 | 5 | 0 | 0 | 105 | 0 | 174 | 210 | 0 | 50 | 5 | 0 |
| Bacteria | Proteobac | Gammaproteo | Pseudomonadale | Moraxellaceae | Psychrobacter | Psychrobacter alimentarius | TACAGAG | Cluster_38 | 729 | 0 | 0 | 108 | 17 | 18 | 1 | 0 | 0 | 16 | 0 | 0 | 0 | 0 | 3 | 552 | 13 | 1 | 0 | 0 |
| Bacteria | Firmicutes | Bacilli | Bacillales | Bacillaceae | Bacillus | Bacillus sp. | TACGTAG | Cluster_39 | 712 | 0 | 0 | 0 | 0 | 0 | 13 | 297 | 355 | 0 | 0 | 0 | 0 | 1 | 0 | 0 | 0 | 25 | 21 | 0 |
| Bacteria | Firmicutes | Bacilli | Lactobacillales | Lactobacillaceae | Lacticaseibacillus | Lacticaseibacillus sp. | TACGTAG | Cluster_40 | 761 | 244 | 0 | 1 | 4 | 2 | 383 | 1 | 0 | 0 | 0 | 0 | 117 | 0 | 0 | 4 | 5 | 0 | 0 | 0 |
| Bacteria | Proteobac | Gammaproteo | Pseudomonadale | Moraxellaceae | Acinetobacter | Acinetobacter sp. | TACAGAG | Cluster_41 | 1037 | 0 | 0 | 75 | 45 | 0 | 1 | 0 | 0 | 22 | 893 | 0 | 0 | 0 | 0 | 0 | 1 | 0 | 0 | 0 |
| Bacteria | Proteobac | Alphaproteoba | Caulobacterales | Caulobacteraceae | Brevundimonas | Brevundimonas sp. | TACGAAG | Cluster_42 | 568 | 0 | 0 | 92 | 10 | 0 | 0 | 2 | 0 | 3 | 460 | 0 | 1 | 0 | 0 | 0 | 0 | 0 | 0 | 0 |
| Bacteria | Firmicutes | Bacilli | Lactobacillales | Lactobacillaceae | Latilactobacillus | Latilactobacillus sp. | TACGTAG | Cluster_43 | 790 | 0 | 0 | 0 | 0 | 0 | 1 | 0 | 0 | 565 | 0 | 0 | 14 | 43 | 0 | 0 | 0 | 0 | 1 | 166 |
| Bacteria | Firmicutes | Bacilli | Lactobacillales | Streptococcaceae | Lactococcus | Lactococcus sp. | TACGTAG | Cluster_44 | 582 | 33 | 80 | 0 | 0 | 0 | 13 | 0 | 0 | 122 | 0 | 242 | 31 | 35 | 0 | 0 | 0 | 0 | 0 | 26 |
| Bacteria | Firmicutes | Bacilli | Lactobacillales | Streptococcaceae | Lactococcus | Lactococcus sp. | TACGTAG | Cluster_45 | 609 | 21 | 2 | 1 | 1 | 2 | 354 | 0 | 0 | 3 | 0 | 1 | 127 | 85 | 1 | 9 | 2 | 0 | 0 | 0 |
| Bacteria | Firmicutes | Bacilli | Lactobacillales | Lactobacillaceae | Latilactobacillus | Latilactobacillus sp. | TACGTAG | Cluster_46 | 619 | 0 | 0 | 0 | 16 | 0 | 0 | 1 | 0 | 296 | 0 | 0 | 0 | 0 | 0 | 0 | 0 | 0 | 0 | 306 |
| Bacteria | Firmicutes | Bacilli | Lactobacillales | Enterococcaceae | Enterococcus | Enterococcus sp. | TACGTAG | Cluster_47 | 677 | 0 | 0 | 0 | 1 | 0 | 9 | 175 | 83 | 1 | 0 | 0 | 66 | 0 | 3 | 0 | 0 | 216 | 123 | 0 |
| Bacteria | Proteobac | Gammaproteo | Pseudomonadale | Pseudomonadaceae | Pseudomonas | Pseudomonas sp. | TACAGAG | Cluster_48 | 621 | 0 | 0 | 27 | 19 | 0 | 429 | 0 | 0 | 3 | 90 | 0 | 27 | 0 | 0 | 0 | 26 | 0 | 0 | 0 |
| Bacteria | Proteobac | Gammaproteo | Pseudomonadale | Moraxellaceae | Enhydrobacter | Moraxella osloensis | TACAGAG | Cluster_49 | 415 | 0 | 0 | 15 | 4 | 0 | 0 | 0 | 0 | 6 | 388 | 2 | 0 | 0 | 0 | 0 | 0 | 0 | 0 | 0 |
| Bacteria | Firmicutes | Bacilli | Lactobacillales | Streptococcaceae | Lactococcus | Lactococcus sp. | TACGTAG | Cluster_50 | 385 | 363 | 3 | 2 | 0 | 4 | 3 | 0 | 0 | 0 | 0 | 0 | 3 | 0 | 0 | 7 | 0 | 0 | 0 | 0 |
| Bacteria | Firmicutes | Bacilli | Lactobacillales | Lactobacillaceae | Levilactobacillus | Levilactobacillus brevis | TACGTAG | Cluster_51 | 433 | 0 | 0 | 32 | 44 | 0 | 0 | 0 | 0 | 40 | 312 | 0 | 0 | 0 | 0 | 0 | 0 | 0 | 1 | 4 |
| Bacteria | Firmicutes | Bacilli | Lactobacillales | Lactobacillaceae | Leuconostoc | Leuconostoc sp. | TACGTAG | Cluster_52 | 456 | 0 | 0 | 0 | 0 | 0 | 1 | 27 | 1 | 58 | 0 | 2 | 200 | 0 | 0 | 0 | 0 | 1 | 3 | 163 |
| Bacteria | Proteobac | Gammaproteo | Burkholderiales | Comamonadaceae | Roseateles | Roseateles saccharophilus | TACGTAG | Cluster_53 | 418 | 0 | 0 | 6 | 189 | 0 | 0 | 0 | 0 | 1 | 222 | 0 | 0 | 0 | 0 | 0 | 0 | 0 | 0 | 0 |
| Bacteria | Firmicutes | Bacilli | Lactobacillales | Lactobacillaceae | Lactobacillus | Lactobacillus gasseri | TACGTAG | Cluster_54 | 542 | 0 | 0 | 0 | 534 | 0 | 1 | 1 | 1 | 0 | 0 | 0 | 1 | 2 | 0 | 0 | 2 | 0 | 0 | 0 |
| Bacteria | Proteobac | Gammaproteo | Enterobacterales | Aeromonadaceae | Aeromonas | Aeromonas sp. | TACGGAG | Cluster_55 | 397 | 0 | 0 | 24 | 1 | 0 | 80 | 0 | 0 | 0 | 0 | 0 | 292 | 0 | 0 | 0 | 0 | 0 | 0 | 0 |
| Bacteria | Firmicutes | Bacilli | Lactobacillales | Lactobacillaceae | Lacticaseibacillus | Lacticaseibacillus casei | TACGTAG | Cluster_56 | 1446 | 135 | 0 | 1 | 3 | 327 | 327 | 0 | 0 | 0 | 0 | 0 | 116 | 0 | 212 | 318 | 6 | 1 | 0 | 0 |
| Bacteria | Firmicutes | Bacilli | Lactobacillales | Lactobacillaceae | Leuconostoc | Leuconostoc mesentereoides | TACGTAT | Cluster_57 | 359 | 0 | 0 | 0 | 0 | 0 | 0 | 182 | 0 | 15 | 0 | 0 | 0 | 0 | 0 | 0 | 0 | 115 | 1 | 46 |
| Bacteria | Proteobac | Gammaproteo | Pseudomonadale | Pseudomonadaceae | Pseudomonas | Pseudomonas sp. | TACAGAG | Cluster_58 | 270 | 0 | 0 | 0 | 10 | 0 | 0 | 0 | 0 | 0 | 0 | 0 | 0 | 0 | 0 | 0 | 260 | 0 | 0 | 0 |
| Bacteria | Proteobac | Gammaproteo | Burkholderiales | Burkholderiaceae | Burkholderia | Burkholderia sp. | TACGTAG | Cluster_59 | 345 | 0 | 0 | 14 | 2 | 0 | 0 | 0 | 0 | 0 | 329 | 0 | 0 | 0 | 0 | 0 | 0 | 0 | 0 | 0 |
| Bacteria | Firmicutes | Bacilli | Lactobacillales | Enterococcaceae | Enterococcus | Enterococcus sp. | TACGTAG | Cluster_60 | 350 | 7 | 0 | 0 | 3 | 64 | 74 | 3 | 1 | 2 | 0 | 0 | 48 | 15 | 56 | 75 | 1 | 0 | 1 | 0 |
| Bacteria | Proteobac | Alphaproteoba | Sphingomonadal | Sphingomonadaceae | Sphingomonas | Sphingomonas sp. | TACGGAG | Cluster_61 | 459 | 0 | 0 | 13 | 26 | 0 | 0 | 0 | 0 | 4 | 416 | 0 | 0 | 0 | 0 | 0 | 0 | 0 | 0 | 0 |
| Bacteria | Actinobac | Actinobacteria | Corynebacteriale | Corynebacteriaceae | Lawsonella | Lawsonella clevelandensis | TACGTAG | Cluster_62 | 240 | 0 | 0 | 148 | 74 | 0 | 0 | 0 | 0 | 16 | 0 | 1 | 0 | 0 | 0 | 0 | 0 | 1 | 0 | 0 |
| Bacteria | Proteobac | Gammaproteo | Xanthomonadale | Xanthomonadaceae | Stenotrophomonas | Stenotrophomonas rhizophila | TACGAAG | Cluster_63 | 267 | 0 | 0 | 3 | 0 | 0 | 0 | 0 | 0 | 0 | 222 | 0 | 42 | 0 | 0 | 0 | 0 | 0 | 0 | 0 |
| Bacteria | Bacteroid | Bacteroidia | Flavobacteriales | Flavobacteriaceae | Marinirhabdus | Marinirhabdus citrea | TACGGAG | Cluster_64 | 301 | 0 | 0 | 0 | 0 | 0 | 0 | 0 | 0 | 0 | 301 | 0 | 0 | 0 | 0 | 0 | 0 | 0 | 0 | 0 |
| Bacteria | Firmicutes | Bacilli | Lactobacillales | Lactobacillaceae | Weissella | Weissella soli | TACGTAT | Cluster_65 | 290 | 0 | 0 | 0 | 0 | 0 | 0 | 82 | 0 | 40 | 0 | 0 | 0 | 1 | 0 | 0 | 0 | 73 | 0 | 94 |
| Bacteria | Actinobac | Actinobacteria | Micrococcales | Micrococcaceae | Micrococcus | Micrococcus luteus | TACGTAG | Cluster_66 | 243 | 0 | 0 | 228 | 10 | 0 | 0 | 0 | 0 | 5 | 0 | 0 | 0 | 0 | 0 | 0 | 0 | 0 | 0 | 0 |
| Bacteria | Firmicutes | Bacilli | Lactobacillales | Lactobacillaceae | Fructilactobacillus | Fructilactobacillus sanfrancisce | TACGTAG | Cluster_67 | 213 | 0 | 0 | 99 | 21 | 0 | 0 | 0 | 0 | 93 | 0 | 0 | 0 | 0 | 0 | 0 | 0 | 0 | 0 | 0 |
| Bacteria | Firmicutes | Bacilli | Staphylococcales | Staphylococcaceae | Staphylococcus | Staphylococcus sp. | TACGTAG | Cluster_68 | 385 | 0 | 6 | 0 | 0 | 0 | 0 | 66 | 29 | 0 | 0 | 4 | 0 | 261 | 0 | 0 | 0 | 6 | 13 | 0 |
| Bacteria | Firmicutes | Bacilli | Lactobacillales | Lactobacillaceae | Lactiplantibacillus | Lactiplantibacillus sp. | TACGTAG | Cluster_69 | 252 | 5 | 0 | 0 | 0 | 0 | 30 | 1 | 0 | 46 | 0 | 0 | 161 | 1 | 0 | 0 | 0 | 0 | 0 | 8 |
| Bacteria | Bacteroid | Bacteroidia | Flavobacteriales | Weeksellaceae | Chryseobacterium | Chryseobacterium sp. | TACGGAG | Cluster_70 | 196 | 0 | 0 | 2 | 0 | 0 | 0 | 0 | 0 | 1 | 193 | 0 | 0 | 0 | 0 | 0 | 0 | 0 | 0 | 0 |
| Bacteria | Firmicutes | Bacilli | Lactobacillales | Lactobacillaceae | Weissella | Weissella sp. | TACGTAT | Cluster_71 | 468 | 0 | 0 | 0 | 0 | 0 | 0 | 207 | 0 | 25 | 0 | 0 | 0 | 0 | 0 | 0 | 0 | 147 | 0 | 89 |
| Bacteria | Firmicutes | Bacilli | Lactobacillales | Enterococcaceae | Enterococcus | Enterococcus sp. | TACGTAG | Cluster_72 | 201 | 0 | 0 | 0 | 0 | 0 | 0 | 70 | 27 | 0 | 0 | 0 | 8 | 0 | 0 | 1 | 0 | 36 | 59 | 0 |
| Bacteria | Firmicutes | Bacilli | Lactobacillales | Streptococcaceae | Lactococcus | Lactococcus sp. | TACGTAG | Cluster_73 | 305 | 55 | 0 | 0 | 0 | 0 | 204 | 0 | 0 | 5 | 0 | 0 | 22 | 13 | 0 | 1 | 0 | 0 | 0 | 5 |
| Bacteria | Proteobac | Gammaproteo | Pseudomonadale | Pseudomonadaceae | Pseudomonas | Pseudomonas sp. | TACGAAG | Cluster_74 | 169 | 0 | 0 | 25 | 0 | 0 | 0 | 0 | 0 | 1 | 143 | 0 | 0 | 0 | 0 | 0 | 0 | 0 | 0 | 0 |
| Bacteria | Actinobac | Actinobacteria | Corynebacteriale | Corynebacteriaceae | Corynebacterium | Corynebacterium sp. | TACGTAG | Cluster_75 | 172 | 0 | 0 | 0 | 5 | 0 | 0 | 0 | 0 | 0 | 167 | 0 | 0 | 0 | 0 | 0 | 0 | 0 | 0 | 0 |
| Bacteria | Firmicutes | Bacilli | Lactobacillales | Lactobacillaceae | Secundilactobacillus | Secundilactobacillus sp. | TACGTAG | Cluster_76 | 324 | 4 | 0 | 1 | 0 | 0 | 0 | 9 | 0 | 166 | 0 | 0 | 14 | 0 | 0 | 0 | 0 | 0 | 0 | 130 |
| Bacteria | Firmicutes | Bacilli | Lactobacillales | Lactobacillaceae | Leuconostoc | Leuconostoc mesenteroides | TACAGAG | Cluster_77 | 243 | 1 | 0 | 0 | 0 | 0 | 0 | 31 | 147 | 0 | 0 | 0 | 2 | 0 | 0 | 0 | 0 | 10 | 52 | 0 |
| Bacteria | Firmicutes | Bacilli | Lactobacillales | Lactobacillaceae | Pediococcus | Pediococcus sp. | TACGTAG | Cluster_78 | 718 | 0 | 0 | 0 | 0 | 0 | 1 | 583 | 13 | 1 | 0 | 0 | 5 | 0 | 0 | 0 | 0 | 81 | 19 | 15 |
| Bacteria | Firmicutes | Bacilli | Lactobacillales | Lactobacillaceae | Lacticaseibacillus | Lacticaseibacillus sp. | TACGTAG | Cluster_79 | 350 | 10 | 0 | 0 | 0 | 207 | 2 | 0 | 0 | 0 | 0 | 0 | 4 | 0 | 12 | 115 | 0 | 0 | 0 | 0 |
| Bacteria | Proteobac | Alphaproteoba | Rhizobiales | Rhizobiaceae | Allorhizobium-Neorh | Agrobacterium fabrum | TACGAAG | Cluster_80 | 137 | 0 | 0 | 13 | 0 | 0 | 0 | 0 | 0 | 6 | 118 | 0 | 0 | 0 | 0 | 0 | 0 | 0 | 0 | 0 |
| Bacteria | Firmicutes | Bacilli | Lactobacillales | Streptococcaceae | Lactococcus | Lactococcus sp. | TACGTAG | Cluster_81 | 586 | 49 | 92 | 0 | 0 | 0 | 18 | 0 | 1 | 1 | 0 | 364 | 8 | 50 | 0 | 0 | 0 | 0 | 0 | 3 |
| Bacteria | Actinobac | Actinobacteria | Micrococcales | Micrococcaceae | Rothia | Rothia kristinae | TACGTAG | Cluster_82 | 109 | 0 | 0 | 0 | 0 | 0 | 0 | 34 | 1 | 0 | 0 | 0 | 0 | 0 | 0 | 0 | 0 | 74 | 0 | 0 |
| Bacteria | Firmicutes | Bacilli | Lactobacillales | Lactobacillaceae | Limosilactobacillus | Limosilactobacillus sp. | TACGTAG | Cluster_83 | 289 | 0 | 0 | 0 | 287 | 0 | 0 | 0 | 0 | 0 | 0 | 0 | 0 | 0 | 0 | 0 | 0 | 0 | 1 | 1 |
| Bacteria | Firmicutes | Bacilli | Lactobacillales | Lactobacillaceae | Lactiplantibacillus | Lactiplantibacillus sp. | TACGTAT | Cluster_84 | 180 | 0 | 0 | 0 | 0 | 0 | 2 | 0 | 0 | 8 | 0 | 0 | 161 | 0 | 0 | 0 | 0 | 0 | 0 | 9 |
| Bacteria | Firmicutes | Bacilli | Lactobacillales | Streptococcaceae | Lactococcus | Lactococcus sp. | TACGTAG | Cluster_85 | 480 | 58 | 55 | 0 | 0 | 0 | 44 | 0 | 0 | 5 | 0 | 198 | 14 | 104 | 0 | 0 | 0 | 0 | 0 | 2 |
| Bacteria | Proteobac | Gammaproteo | Pseudomonadale | Halomonadaceae | Halomonas | Halomonas profundi | TACGGAG | Cluster_86 | 196 | 0 | 0 | 124 | 43 | 0 | 1 | 0 | 0 | 9 | 0 | 0 | 0 | 0 | 0 | 0 | 19 | 0 | 0 | 0 |
| Bacteria | Proteobac | Gammaproteo | Enterobacterales | Pseudoalteromonada | Pseudoalteromonas | Pseudoalteromonas sp. | TACGGAG | Cluster_87 | 106 | 0 | 0 | 0 | 0 | 1 | 21 | 0 | 0 | 4 | 0 | 0 | 0 | 0 | 0 | 0 | 80 | 0 | 0 | 0 |
| Bacteria | Proteobac | Gammaproteo | Burkholderiales | Burkholderiaceae | Ralstonia | Ralstonia pickettii | TACGTAG | Cluster_88 | 121 | 0 | 0 | 2 | 118 | 0 | 0 | 0 | 0 | 0 | 0 | 0 | 0 | 0 | 0 | 1 | 0 | 0 | 0 | 0 |
| Bacteria | Actinobac | Actinobacteria | Corynebacteriale | Corynebacteriaceae | Corynebacterium | Corynebacterium sp. | TACGTAG | Cluster_89 | 191 | 0 | 0 | 108 | 78 | 0 | 0 | 0 | 0 | 5 | 0 | 0 | 0 | 0 | 0 | 0 | 0 | 0 | 0 | 0 |
| Bacteria | Actinobac | Actinobacteria | Pseudonocardial | Pseudonocardiaceae | Pseudonocardia | Pseudonocardia sp. | TACGTAG | Cluster_90 | 88 | 0 | 0 | 0 | 0 | 0 | 0 | 0 | 0 | 0 | 88 | 0 | 0 | 0 | 0 | 0 | 0 | 0 | 0 | 0 |
| Bacteria | Firmicutes | Bacilli | Lactobacillales | Streptococcaceae | Lactococcus | Lactococcus sp. | TACGTAG | Cluster_91 | 207 | 0 | 9 | 0 | 0 | 0 | 5 | 0 | 0 | 0 | 0 | 5 | 0 | 188 | 0 | 0 | 0 | 0 | 0 | 0 |
| Bacteria | Firmicutes | Bacilli | Lactobacillales | Enterococcaceae | Enterococcus | Enterococcus sp. | TACGTAT | Cluster_92 | 215 | 0 | 0 | 0 | 0 | 0 | 0 | 52 | 21 | 0 | 0 | 0 | 4 | 0 | 0 | 0 | 0 | 43 | 95 | 0 |
| Bacteria | Firmicutes | Bacilli | Lactobacillales | Enterococcaceae | Enterococcus | Enterococcus sp. | TACGTAG | Cluster_93 | 128 | 0 | 0 | 0 | 0 | 10 | 3 | 44 | 0 | 0 | 0 | 0 | 4 | 0 | 39 | 25 | 0 | 3 | 0 | 0 |
| Bacteria | Proteobac | Gammaproteo | Enterobacterales | Pasteurellaceae | Actinobacillus | Actinobacillus minor | TACGGAG | Cluster_94 | 121 | 0 | 0 | 0 | 121 | 0 | 0 | 0 | 0 | 0 | 0 | 0 | 0 | 0 | 0 | 0 | 0 | 0 | 0 | 0 |
| Bacteria | Firmicutes | Bacilli | Lactobacillales | Enterococcaceae | Enterococcus | Enterococcus sp. | TACGTAG | Cluster_95 | 140 | 0 | 0 | 0 | 0 | 0 | 0 | 53 | 49 | 0 | 0 | 0 | 4 | 4 | 1 | 0 | 0 | 14 | 15 | 0 |
| Bacteria | Bacteroid | Bacteroidia | Bacteroidales | Prevotellaceae | Segatella | Segatella copri | TACGGAA | Cluster_96 | 96 | 0 | 0 | 0 | 96 | 0 | 0 | 0 | 0 | 0 | 0 | 0 | 0 | 0 | 0 | 0 | 0 | 0 | 0 | 0 |
| Bacteria | Firmicutes | Bacilli | Lactobacillales | Enterococcaceae | Enterococcus | Enterococcus sp. | TACGTAT | Cluster_97 | 151 | 0 | 0 | 0 | 0 | 0 | 0 | 35 | 15 | 0 | 0 | 0 | 4 | 0 | 0 | 0 | 1 | 17 | 79 | 0 |
| Bacteria | Firmicutes | Bacilli | Lactobacillales | Lactobacillaceae | Weissella | Weissella sp. | TACGTAT | Cluster_98 | 225 | 0 | 0 | 0 | 0 | 0 | 0 | 65 | 0 | 91 | 0 | 0 | 0 | 10 | 0 | 0 | 0 | 17 | 0 | 42 |
| Bacteria | Firmicutes | Bacilli | Lactobacillales | Enterococcaceae | Enterococcus | Enterococcus sp. | TACGTAG | Cluster_99 | 72 | 0 | 0 | 0 | 0 | 19 | 0 | 0 | 0 | 0 | 0 | 0 | 0 | 0 | 33 | 20 | 0 | 0 | 0 | 0 |
| Bacteria | Firmicutes | Bacilli | Lactobacillales | Lactobacillaceae | Lacticaseibacillus | Lacticaseibacillus sp. | TACGTAT | Cluster_100 | 62 | 0 | 0 | 0 | 0 | 4 | 0 | 51 | 0 | 0 | 0 | 0 | 1 | 0 | 0 | 0 | 0 | 5 | 1 | 0 |
| Bacteria | Firmicutes | Bacilli | Lactobacillales | Enterococcaceae | Enterococcus | Enterococcus sp. | TACGTAG | Cluster_101 | 65 | 0 | 0 | 0 | 0 | 0 | 0 | 29 | 8 | 0 | 0 | 0 | 0 | 1 | 1 | 0 | 0 | 12 | 14 | 0 |
| Bacteria | Firmicutes | Bacilli | Mycoplasmatale | Mycoplasmataceae | Mesomycoplasma | Mesomycoplasma hyopneumon | TACATAG | Cluster_102 | 77 | 0 | 0 | 0 | 77 | 0 | 0 | 0 | 0 | 0 | 0 | 0 | 0 | 0 | 0 | 0 | 0 | 0 | 0 | 0 |
| Bacteria | Firmicutes | Bacilli | Lactobacillales | Lactobacillaceae | Leuconostoc | Leuconostoc sp. | TACGTAT | Cluster_103 | 225 | 1 | 0 | 0 | 0 | 0 | 0 | 106 | 24 | 0 | 0 | 1 | 6 | 0 | 0 | 0 | 0 | 25 | 62 | 0 |
| Bacteria | Firmicutes | Bacilli | Lactobacillales | Lactobacillaceae | Weissella | Weissella soli | TACGTAT | Cluster_104 | 115 | 0 | 0 | 0 | 0 | 0 | 0 | 42 | 0 | 7 | 0 | 0 | 0 | 0 | 0 | 0 | 0 | 23 | 0 | 43 |
| Bacteria | Firmicutes | Bacilli | Lactobacillales | Lactobacillaceae | Lacticaseibacillus | Lacticaseibacillus sp. | TACGTAG | Cluster_105 | 137 | 0 | 0 | 0 | 0 | 31 | 0 | 0 | 0 | 0 | 0 | 0 | 0 | 0 | 40 | 66 | 0 | 0 | 0 | 0 |
| Bacteria | Firmicutes | Bacilli | Lactobacillales | Enterococcaceae | Enterococcus | Enterococcus sp. | TACGTAG | Cluster_106 | 131 | 0 | 0 | 0 | 0 | 21 | 0 | 0 | 0 | 0 | 0 | 0 | 0 | 0 | 62 | 48 | 0 | 0 | 0 | 0 |
| Bacteria | Actinobac | Actinobacteria | Bifidobacteriales | Bifidobacteriaceae | Bifidobacterium | Bifidobacterium sp. | TACGTAG | Cluster_107 | 78 | 0 | 0 | 0 | 77 | 1 | 0 | 0 | 0 | 0 | 0 | 0 | 0 | 0 | 0 | 0 | 0 | 0 | 0 | 0 |
| Bacteria | Firmicutes | Bacilli | Lactobacillales | Lactobacillaceae | Agrilactobacillus | Agrilactobacillus sp. | TACGTAG | Cluster_108 | 64 | 0 | 0 | 0 | 0 | 0 | 3 | 0 | 0 | 37 | 0 | 0 | 11 | 2 | 0 | 0 | 0 | 0 | 0 | 11 |
| Bacteria | Firmicutes | Bacilli | Lactobacillales | Lactobacillaceae | Leuconostoc | Leuconostoc mesenteroides | TACGTAT | Cluster_109 | 85 | 1 | 0 | 0 | 0 | 0 | 0 | 20 | 10 | 0 | 0 | 0 | 0 | 0 | 0 | 0 | 0 | 15 | 39 | 0 |
| Bacteria | Firmicutes | Bacilli | Lactobacillales | Lactobacillaceae | Latilactobacillus | Latilactobacillus sp. | TACGTAG | Cluster_110 | 76 | 0 | 0 | 0 | 0 | 0 | 0 | 0 | 0 | 5 | 0 | 0 | 70 | 0 | 0 | 0 | 0 | 0 | 0 | 1 |
| Bacteria | Firmicutes | Bacilli | Lactobacillales | Lactobacillaceae | Lactobacillus | Lactobacillus sp. | TACGTAG | Cluster_111 | 212 | 0 | 0 | 0 | 0 | 44 | 0 | 0 | 0 | 0 | 0 | 0 | 0 | 0 | 91 | 77 | 0 | 0 | 0 | 0 |
| Bacteria | Firmicutes | Bacilli | Lactobacillales | Lactobacillaceae | Lacticaseibacillus | Lacticaseibacillus sp. | TACGTAG | Cluster_112 | 94 | 0 | 0 | 0 | 0 | 30 | 0 | 0 | 0 | 0 | 0 | 0 | 0 | 0 | 23 | 41 | 0 | 0 | 0 | 0 |
| Bacteria | Firmicutes | Bacilli | Bacillales | Bacillaceae | Bacillus | Bacillus sp. | TACGTAG | Cluster_113 | 75 | 0 | 0 | 0 | 0 | 0 | 0 | 23 | 46 | 0 | 0 | 0 | 0 | 0 | 0 | 0 | 0 | 4 | 2 | 0 |
| Bacteria | Firmicutes | Bacilli | Lactobacillales | Enterococcaceae | Enterococcus | Enterococcus sp. | TACGTAG | Cluster_114 | 59 | 0 | 0 | 0 | 0 | 0 | 0 | 23 | 0 | 0 | 0 | 0 | 33 | 0 | 1 | 0 | 0 | 1 | 1 | 0 |
| Bacteria | Firmicutes | Bacilli | Lactobacillales | Streptococcaceae | Lactococcus | Lactococcus sp. | TACGTAG | Cluster_115 | 76 | 7 | 0 | 0 | 1 | 0 | 13 | 0 | 0 | 11 | 0 | 0 | 21 | 0 | 0 | 3 | 0 | 0 | 0 | 20 |
