## Supplementary material for "Ripened plant-based cheese analogs in Europe: nutritional and microbial profiles": Table S4 Fungi

| Kingdom | Phylum | Class | Oder | Family | Genus | Species | seed_sequ | observation_ | observatic | R1 | R2 | R3 | R4 | R5 | R6 | R7 | R8 | R10 |
|---|---|---|---|---|---|---|---|---|---|---|---|---|---|---|---|---|---|---|
| k__Fungi | p__Ascomycota | c__Saccharomycetes | o__Saccharomycetale | f__Dipodascaceae | g__Geotrichum | s__Geotrichum candidum | AAGGATC | Cluster_1 | 319573 | 101 | 65595 | 3178 | 91195 | 40 | 25 | 99899 | 15 | 59525 |
| k__Fungi | p__Ascomycota | c__Saccharomycetes | o__Saccharomycetale | f__Dipodascaceae | g__Geotrichum | s__Geotrichum candidum | AAGGATC | Cluster_2 | 114959 | 1145 | 844 | 109617 | 1628 | 17 | 9 | 1661 | 8 | 30 |
| k__Fungi | p__Ascomycota | c__Saccharomycetes | o__Saccharomycetale | f__Dipodascaceae | g__Dipodascus | s__Geotrichum candidum | AAGGATC | Cluster_3 | 34776 | 4 | 2 | 3 | 24424 | 13 | 2 | 5 | 3 | 10320 |
| k__Fungi | p__Ascomycota | c__Saccharomycetes | o__Saccharomycetale | f__Dipodascaceae | g__Geotrichum | s__Geotrichum candidum | AAGGATC | Cluster_4 | 40665 | 1 | 14465 | 345 | 65 | 1 | 1 | 20098 | 2 | 5687 |
| k__Fungi | p__Ascomycota | c__Saccharomycetes | o__Saccharomycetale | f__Debaryomycetaceae | g__Debaryomyces | s__Debaryomyces hansenii | AAGGATC | Cluster_5 | 42330 | 36151 | 5857 | 239 | 28 | 17 | 10 | 26 | 2 | 0 |
| k__Fungi | p__Ascomycota | c__Eurotiomycetes | o__Eurotiales | f__Aspergillaceae | g__Penicillium | s__Penicillium camemberti | AAGGATC | Cluster_6 | 26923 | 2505 | 4 | 22 | 6 | 12628 | 11413 | 0 | 0 | 345 |
| k__Fungi | p__Ascomycota | c__Saccharomycetes | o__Saccharomycetale | f__Dipodascaceae | g__Geotrichum | s__Geotrichum candidum | AAGGATC | Cluster_7 | 19121 | 5 | 1 | 1 | 3 | 6 | 6 | 5 | 1 | 19093 |
| k__Fungi | p__Ascomycota | c__Saccharomycetes | o__Saccharomycetale | f__Dipodascaceae | g__Geotrichum | s__Geotrichum candidum | AAGGATC | Cluster_8 | 14622 | 0 | 1 | 1 | 1 | 0 | 1 | 3 | 4 | 14611 |
| k__Fungi | p__Ascomycota | c__Eurotiomycetes | o__Eurotiales | f__Aspergillaceae | g__Penicillium | s__Penicillium roqueforti | AAGGATC | Cluster_9 | 12401 | 0 | 0 | 0 | 0 | 0 | 0 | 0 | 12401 | 0 |
| k__Fungi | p__Ascomycota | c__Saccharomycetes | o__Saccharomycetale | f__Saccharomycetacea | g__Torulaspora | s__Torulaspora delbrueckii | AAGGATC | Cluster_10 | 3048 | 0 | 3048 | 0 | 0 | 0 | 0 | 0 | 0 | 0 |
| k__Fungi | p__Ascomycota | c__Saccharomycetes | o__Saccharomycetale | f__Dipodascaceae | g__Geotrichum | s__Geotrichum candidum | AAGGATC | Cluster_11 | 142 | 0 | 0 | 1 | 0 | 0 | 0 | 0 | 0 | 141 |
| k__Fungi | p__Ascomycota | c__Saccharomycetes | o__Saccharomycetale | f__Metschnikowiaceae | g__Clavispora | s__Clavispora lusitaniae | AAGGATC | Cluster_12 | 46 | 11 | 30 | 4 | 1 | 0 | 0 | 0 | 0 | 0 |
