## Supplementary material for "Ripened plant-based cheese analogs in Europe: nutritional and microbial profiles": Table S1

| Sample | Geographical origin | Producer | Ingredients | Rind/ Technology | Photo |
|---|---|---|---|---|---|
| **C1** | Ile-de-France, France | P1 | water, cashew nuts, salt, starter cultures | Bloomy, soft curd | 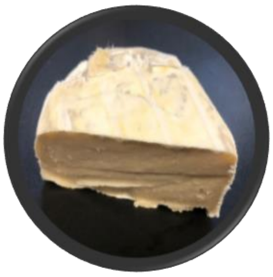 |
| **C2** | | | water, cashew nuts, salt, starter cultures | Natural, soft curd | 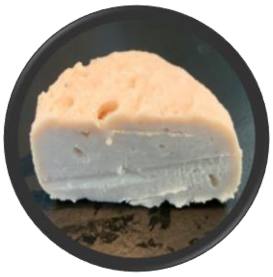 |
| **C3** | | | water, cashew nuts, soy, salt, starter cultures | Bloomy, soft curd | 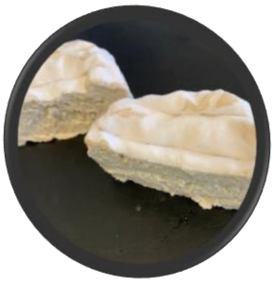 |
| **C4** | Ile-de-France, France | P2 | cashew nuts, pumpkin seeds, pine nuts, water and salt | Bloomy, soft curd | 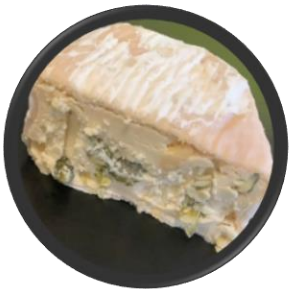 |
| **C5** | Amsterdam, The Netherlands | P3 | cashew nuts, water, black summer truffle with naturally extra virgin olive oil, vegan *P. camemberti* (0.003%) | Bloomy, soft curd | 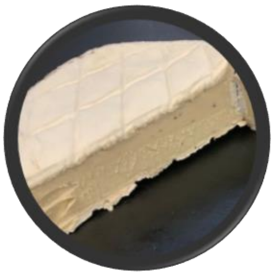 |

| | | | | | |
|---|---|---|---|---|---|
| **C6** | | | cashew nuts, water, salt, pepper, black, vegetal actif charcoal, vegan starter culture *P. camemberti* (0.003%) | Bloomy, soft curd | 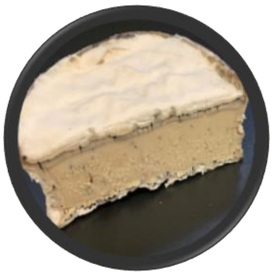 |
| **C7** | Ile-de-France, France | P4 | cashew nuts, water, salt, black pepper, vegetable charcoal, vegan starter culture, urucum seed, ferments | Washed, soft curd | 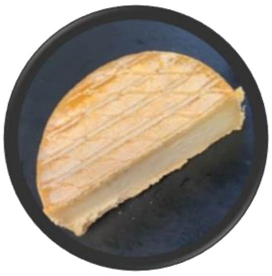 |
| **C8** | | P5 | cashew nuts, water, coconut milk, salt, ferments. | Natural, blue curd | 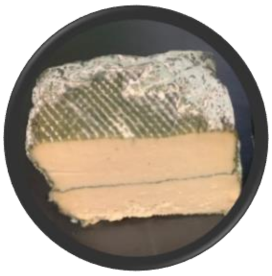 |
| **C9** | | | cashew nuts, water, salt, ferments. | Bloomy, soft curd | 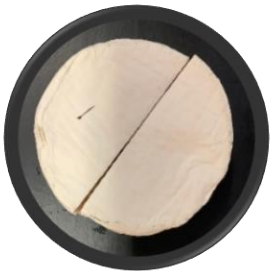 |
| **C10** | | P6 | organic fresh almond milk, cashew nuts, salt ferments | Bloomy, soft curd | 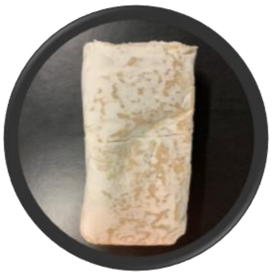 |
| **C11** | | | organic fresh almond milk, cashew nuts, salt ferments | Bloomy, soft curd | 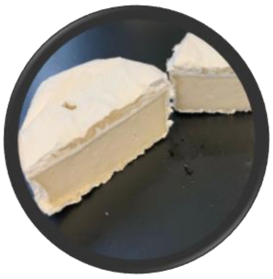 |
